# Cytoplasmic import and processing of mRNA amplify transcriptional bursts accounting for the majority of cellular noise

**DOI:** 10.1101/222901

**Authors:** Maike M. K. Hansen, Ravi V. Desai, Michael L. Simpson, Leor S. Weinberger

## Abstract

Transcription is an episodic process characterized by probabilistic bursts; but how these bursts are modulated by cellular physiology remains unclear and has implications for how selection may act on these fluctuations. Using simulations and single-molecule RNA counting, we examined how cellular processes influence cell-to-cell variability (noise). The results show that RNA noise is amplified in the cytoplasm compared to the nucleus in ~85% of genes across diverse promoters, genomic loci, and cell types (human and mouse). Surprisingly, measurements show further amplification of RNA noise in the cytoplasm, fitting a model of biphasic mRNA conversion between translation- and degradation-competent states. The multi-state translation-degradation of mRNA also causes substantial noise amplification in protein levels, ultimately accounting for ~74% of intrinsic protein variability in cell populations. Overall, the results demonstrate how transcriptional bursts are intrinsically amplified by mRNA processing and indicate mechanisms through which noise could act as a substrate for evolutionary selection.

## Introduction

Intracellular biological reactions can exhibit large intrinsic fluctuations (i.e., stochastic ‘noise’) that manifest as cell-to-cell variability, even in isogenic populations of cells (Blake et al., 2003; Elowitz et al., 2002; Kaern et al., 2005; Kepler and Elston, 2001). These intrinsic stochastic fluctuations partly originate during transcription (Golding et al., 2005; Raj et al., 2006) and drive strong evolutionary selection pressures (Fraser et al., 2004; Metzger et al., 2015) as well as cell-fate decisions (Balázsi et al., 2011; Suel et al., 2007; Weinberger et al., 2005).

Transcriptional fluctuations can be largely due to the episodic nature of transcription, commonly called ‘bursting’, in which short periods of productive promoter activity are interspersed between long periods of promoter inactivity (Chong et al., 2014; Coulon et al., 2013; Dar et al., 2012; Golding et al., 2005; Raj et al., 2006; Sanchez and Golding, 2013; Singh et al., 2010; Suter et al., 2011; Zenklusen et al., 2008). These episodic transcriptional bursts appear to be predominant in mammalian cells, especially at low transcript abundance (Dar et al., 2012; Raj et al., 2006). In the common transcriptional bursting model, the ‘two-state random-telegraph’ model, a promoter toggles between a transcriptionally inactive OFF state and an active ON state (Kepler and Elston, 2001). While more than two promoter states may exist, all multistate transcription models generate super-Poissonian cell-to-cell distributions (noise) in mRNA and protein, especially for the relatively slow toggling rates measured for many promoters (Harper et al., 2011; Zenklusen et al., 2008). These transcriptional bursting models contrast with minimally stochastic, single-state (i.e., constitutive) transcription models, which are Poisson processes and generate Poisson distributions for cell-to-cell variability in gene products. These Poisson distributions represent the theoretical low-noise limit for gene expression (Kaern et al., 2005), but the more complex multi-state models (e.g. random-telegraph models) are required to fit the vast majority of measured cell-to-cell expression distributions, which are super-Poissonian (Sanchez and Golding, 2013). While variability in the abundance of nascent transcripts—those still tethered to DNA at the transcriptional center—can fall under the Poisson limit (Choubey et al., 2015), this scenario is not a birth-death process but a special case of an age-structured process described by a particular form of the gamma distribution called an Erlang distribution (Mittler et al., 1998). Nevertheless, once transcripts are released from the DNA, the process can no longer be considered age structured and the distributions are, at best, Poisson birth-death limited.

Noise that originates during transcription can be modulated by various cellular mechanisms. For example, translation often amplifies transcriptional bursting noise (Ozbudak et al., 2002) and auto-regulatory gene circuits can, depending upon their architecture, either amplify or attenuate noise for their specific target genes (Arias and Hayward, 2006; Austin et al., 2006; Barkai and Leibler, 2000; Isaacs et al., 2003). However, recent studies have suggested that transcriptional noise is efficiently and non-specifically buffered to minimal Poisson levels by ‘passive’ cellular compartmentalization, specifically nuclear export (Battich et al., 2015; Stoeger et al., 2016). The resulting conundrum is, if compartmentalization broadly buffers noise to minimal levels, why is the signature of evolutionary selection visible on promoter architecture? Moreover, how can noise drive selection pressures and cell fate decisions? For example, how are transcriptional regulatory circuits able to modulate noise—i.e., attenuate (Arias and Hayward, 2006) or amplify (Weinberger et al., 2005) noise—when nuclear export acts as a strong downstream filter reducing noise to the theoretical limit?

Given the literature reporting super-Poissonian cytoplasmic mRNA and protein noise in nucleated eukaryotic cells, we sought to reconcile how evidence of super-Poissonian noise could co-exist with nuclear export passively buffering noise to minimal levels. Using computational modeling and single-molecule quantitation by RNA Fluorescence in situ Hybridization (FISH), we predict and then experimentally measure mRNA noise in the nucleus and cytoplasm. We model across the known physiological parameter range and find that in the vast majority of cases (~85%), mRNA noise is amplified by export from the nucleus and is super-Poissonian in the cytoplasm. smFISH measurements corroborate this finding for diverse promoters (LTR, UBC, Ef1-1α, SV40, c-Jun, c-Fos, COX-2, FoxO, Per1, NR4A2 and NANOG) in different cell types. As predicted by modeling, modulation of nuclear export has little effect in changing this amplification. Surprisingly, the smFISH measurements and perturbation experiments indicate a further post-export step of noise-amplification for cytoplasmic mRNA, which supports mRNA translation and degradation being mutually exclusive (multi-state) processes. Finally, we present a model that quantifies how mRNAs are amplified during progression from transcription to translation and predicts super-Poissonian protein noise from transcriptional measures. Overall, the findings demonstrate that transcriptional noise is intrinsically amplified in cells, showing how, in principle, noise could have served as a substrate for promoter selection and can act as a driving force in cell-fate decisions.

## Results

### The standard model of gene expression predicts that—in the physiological parameter regime—mRNA noise is often amplified in the cytoplasm compared to the nucleus

To explore how cellular physiology influences gene expression noise, we used Gillespie’s method (Gillespie, 1977) to perform stochastic simulations of a conventional model of eukaryotic mRNA transcription (Dar et al., 2014; Raj et al., 2006; Raser and O’Shea, 2004; Suter et al., 2011), expanded to include both the nuclear and cytoplasmic compartments (Figure 1A). Since most genes are co-transcriptionally spliced (Tilgner et al., 2012), splicing was incorporated into the transcription rate. A total of 7776 (6^5^) parameter combinations were examined—with 1000 simulations run per parameter combination (i.e., over 7 million simulation runs)—allowing us to vary the rate of each cellular process (e.g., transcription, export, decay) over several orders of magnitude based on literature estimates (Bahar Halpern et al., 2015a; Bahar Halpern et al., 2015b; Battich et al., 2015; Dar et al., 2012; Harper et al., 2011; Suter et al., 2011). Mean (μ) and variance (σ^2^) in mRNA counts were determined for both nuclear and cytoplasmic compartments (Figures S1A).

**Figure 1:**
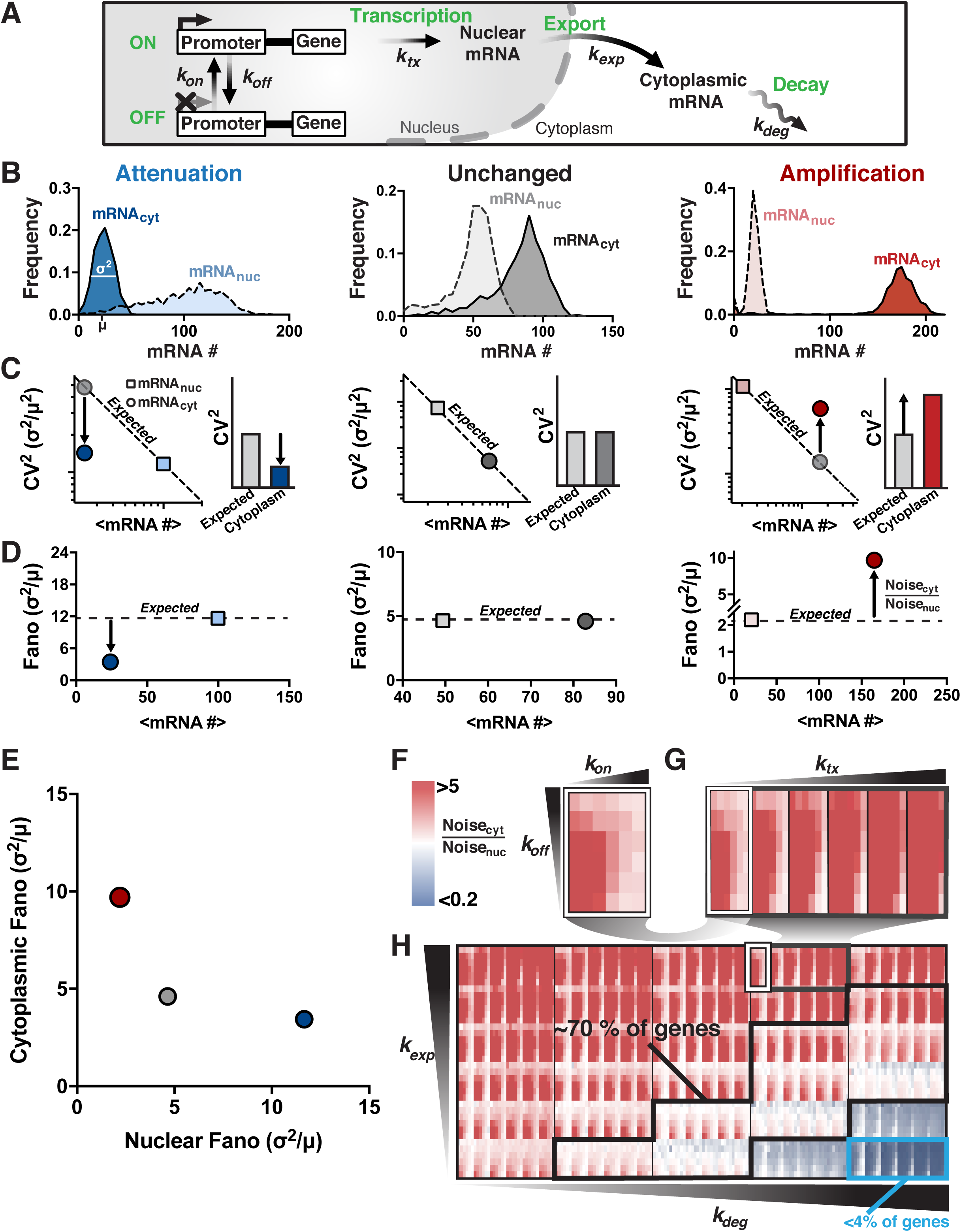
The random-telegraph model of gene expression predicts that mRNA noise is amplified by nuclear-to-cytoplasmic export. (**A**) Schematic representation of the conventional model of eukaryotic mRNA transcription expanded to include both nuclear and cytoplasmic compartments. (**B**) Representative distributions of nuclear (dashed lines) and cytoplasmic (solid lines) mRNA for parameter combinations yielding noise attenuation (blue), unchanged noise (black), and noise amplification (red). Distributions are from 1000 simulations per parameter condition. (**C**) Mean versus CV^2^ for nuclear (squares), expected cytoplasmic (grey circles) and cytoplasmic (colored circles) mRNAs, corresponding to each distribution in B. Bar graphs show the expected cytoplasmic CV^2^ due to Poisson scaling and the actual cytoplasmic CV^2^. (**D**) Mean versus σ^2^/μ (Fano factor) for both nuclear (squares) and cytoplasmic (circles) mRNAs, corresponding to each distribution in B. (**E**) Comparison of nuclear versus cytoplasmic mRNA noise (σ^2^/μ). (**F-H**) Nuclear-to-cytoplasmic noise ratio (Noise_cyt_ /Noise_nuc_) simulated for the physiologically *possible* parameter space, as calculated by varying each parameter from its highest-reported to its lowest-reported value (1000 simulations run per parameter combination; > 7 million runs). Increasing red represents increasing noise amplification while increasing blue represents increasing noise attenuation, white represents no change in noise from nucleus to cytoplasm. Panel F (a subpanel of G) shows how varying *k_on_* and *k_off_* across the full range of reported values, affects the noise ratio (all other parameters are kept fixed). Panel G (a subpanel of H) shows how varying *k_tx_* across its full range of reported values affects the noise ratio for the array of *k_on_ k_off_* simulations. Panel H represents the full set of simulation results where the array of *k_on_ k_off_ k_tx_* simulations is varied over the full reported range of *k_exp_* and *k_deg_* values. The *probable* parameter space (70% of measurements) is marked by the black box, whereas the cyan box (< 4% of measurements) represents the regime of efficient buffering.

When comparing mRNA noise in the nucleus and cytoplasm, three scenarios are possible: (i) Noise can be lower in the cytoplasm than in the nucleus (i.e. *attenuated*) (Figure 1B, blue); (ii) noise can be the same in both compartments (i.e. *unchanged*) (Figure 1B, grey); or (iii), noise can be higher in the cytoplasm than the nucleus (i.e. *amplified*) (Figure 1B, red). When interpreting the coefficient of variation (CV^2^ = σ^2^/μ^2^) for these three scenarios, it is important to note that a decrease in CV^2^ does not necessarily translate to a decrease in noise (Figure 1C). Any changes in rates which cause an increase in mean without an increase in noise will follow so called “Poisson-scaling”—CV^2^ *decreases* to the same extent that the mean *increases* (Kaern et al., 2005). In other words, a decrease in CV^2^ is only an effective attenuation of noise if the decrease is greater than the decrease that would be obtained by a simple scaling of the mean (i.e., attenuated noise only occurs when the CV^2^ falls below the “expected” dashed diagonal in Figure 1C, left). Whereas, noise is effectively “unchanged” when CV^2^ scales as expected from the mean (Figure 1C, middle) and noise is effectively amplified when CV^2^ is higher than expected given the mean (Figure 1C, right). Since these scaling properties can make the CV^2^ a somewhat complex metric, noise is often calculated by variance over the mean (σ^2^/μ; a.k.a, the Fano factor) (Thattai and van Oudenaarden, 2001), where the expected noise is constant with respect to the mean such that attenuation and amplification are readily apparent (Figure 1D-E). As such, the Fano factor automatically provides a measure of deviation from a Poisson process, where σ^2^/μ = 1. The respective noise ratio—Noise_cytoplasm_ / Noise_nucleus_ (which is equivalent to CV^2^_cytoplasm_ /CV^2^_expected_)—was examined for all 7776 parameter combinations (Figure 1F-H).

Remarkably, the results show that for most combinations of physiologically relevant parameters, mRNA noise is largely amplified in the cytoplasm compared to the nucleus (Figure 1F–H, red rectangles). Moreover, the *possible* physiological parameter space can be further limited to a *probable* regime using previously reported genome-wide mRNA counts (Bahar Halpern et al., 2015a). Namely, the reported nuclear and cytoplasmic mRNA counts were used to estimate likely ratios of mRNA export-to-degradation rates (Figure S1C, and Methods Equations 1–5), which largely determine whether noise is amplified, unchanged, or attenuated. This data constraint is applied to generate a *probable* physiological parameter regime in which amplification becomes even more prevalent (Figure 1H, black box). Specifically, about 15% of genes across the genome show >20-fold higher export rates than degradation rates, thus falling within the parameter regime of highly amplified cytoplasmic noise. Another 70% of genes across the genome have significantly faster rates of export than degradation, also falling in the parameter regime of amplification. Finally, only ~15% of genes across the genome fall in the parameter regime in which the rate of export is slower than cytoplasmic mRNA degradation, of which less than 4% have rates where substantial noise attenuation (>5-fold) is even possible (Figure 1H, light blue box). Thus, the data constraints show that ~85% of genes fall in the parameter regime in which noise is amplified in the cytoplasm and only about 2.5% of genes fall in the parameter regime where noise is attenuated down to minimally stochastic Poisson levels—substantially less than previously implied (Battich et al., 2015).

Analytically, a fairly simply expression for the Fano factor ratio between cytoplasm and nucleus can be obtained (see Methods: Analytical derivation):

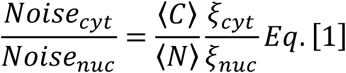

Where 〈*N*〉 and 〈*C*〉 are the mean mRNA abundances in the nucleus and cytoplasm, respectively, while *ξ_cyt_* and *ξ_nuc_* are the noise bandwidths (Simpson et al., 2003) in the cytoplasm and nucleus, respectively. In both cases, the noise bandwidth is dominated by the lowest critical frequency it is associated with (i.e. either the critical frequency of promoter toggling or mRNA export for *ξ_nuc_*, and either the critical frequency of promoter toggling, mRNA export or degradation for *ξ_cyt_*). Intuitively, this means that *ξ_nuc_*≥ *ξ_cyt_*, since *ξ_cyt_* can be dominated by the additional critical frequency associated with degradation, which has no impact on *ξ_nuc_*. Therefore, for all cases Eq. [1] reduces to 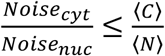, which predicts that there is a strong tendency for *Noise_cyt_* > *Noise_nuc_* when 〈*C*〉 > 〈*N*〉. Given previous reports that most genes exhibit 〈*C*〉 > 〈*N*〉 (Bahar Halpern et al., 2015), most genes are expected to fall in the amplification regime as the numerical simulations show (Figure 1H).

### Single-molecule mRNA quantification shows generalized amplification of noise in the cytoplasm

To experimentally test the model predictions that noise is generally amplified in the cytoplasm, we used single-molecule RNA Fluorescence in situ Hybridization (smFISH) to quantify individual mRNA transcripts in both the nucleus and cytoplasm. To span across the physiological parameter regime, we examined both a panel of GFP-expressing reporter constructs with different promoter architectures, which exhibit widely different transcriptional bursting and expression rates (Figure 2), as well as endogenous genes c-Jun, c-Fos, COX-2, PER1, FoxO, NR4A2 and NANOG (Figure 3).

**Figure 2:**
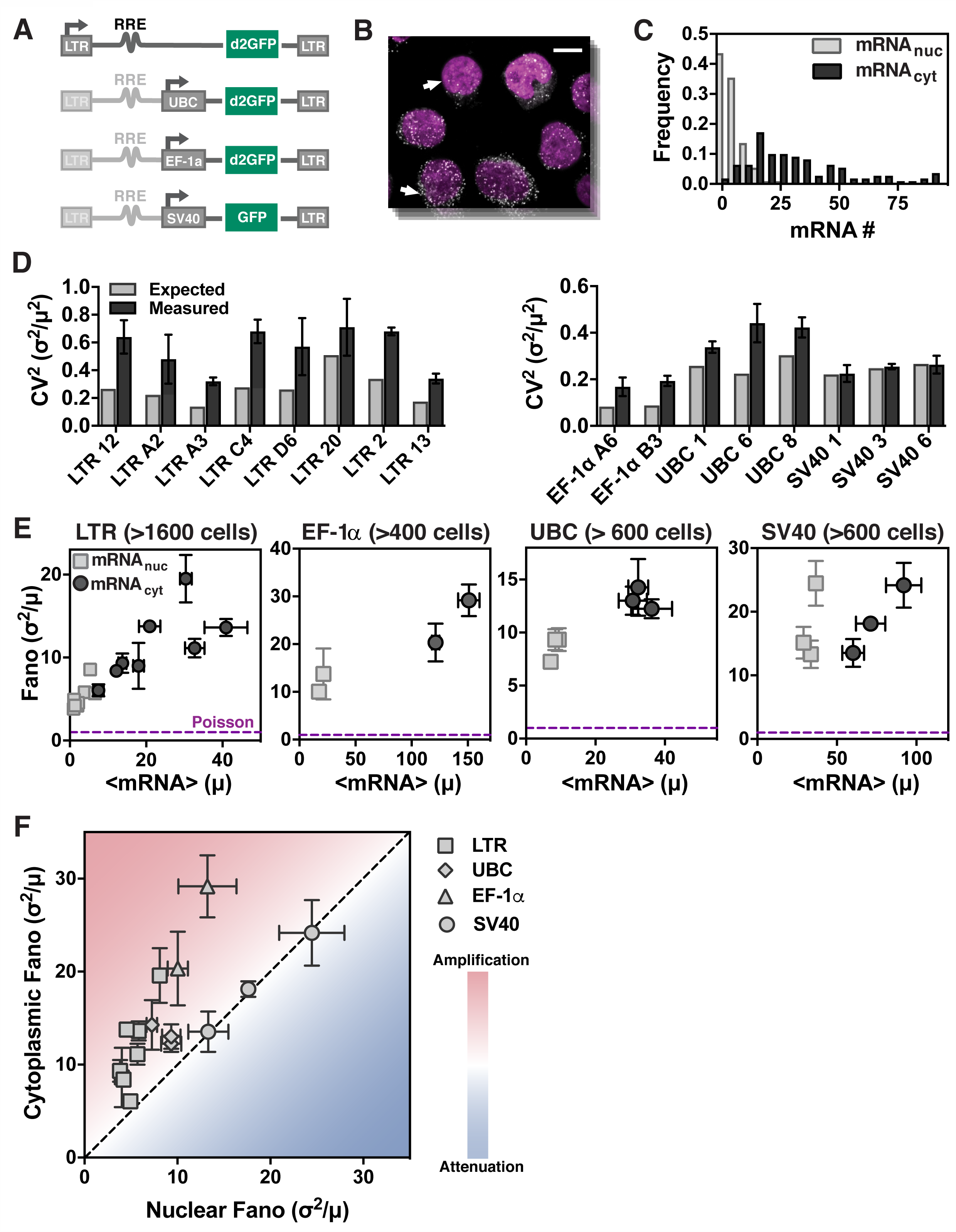
Single-molecule mRNA counting shows amplification of noise in the cytoplasm, independent of promoter type and genomic locus. (**A**) Schematic of reporter constructs used to express mRNAs from the HIV-1 LTR promoter in isoclonal populations of human T lymphocytes (Jurkat) and from the UBC, SV40, and EF-1α promoters in isoclonal populations of human myeloid leukemia cells (K562). (**B**) Representative smRNA FISH micrograph of an isoclonal T-lymphocyte population (maximum intensity projection of 15 optical sections, each spaced 0.4 μm apart) where DNA has been DAPI stained (purple) and mRNAs (white dots) are GFP mRNAs expressed from a single lentiviral-vector integration of a GFP-reporter cassette. Scale bar represents 5 μm, and arrows point towards two similarly sized cells that show high variability in mRNA levels. (**C**) Typical probability distribution of cytoplasmic and nuclear mRNA numbers for a single mRNA reporter species (e.g. GFP mRNA) in an isoclonal population of T cells, after extrinsic noise filtering. (**D**) Expected (grey) versus measured (from smFISH, black) cytoplasmic CV^2^ of mRNAs expressed from all four promoters. (**E**) Mean mRNA expression (μ) versus noise (σ^2^/μ) for both nuclear (squares) and cytoplasmic (circles) mRNAs. Data points are biological replicates, and error bars represent SEM. The minimal noise defined by a Poisson process is shown as a purple line (σ^2^/μ = 1). (**F**) Comparison of nuclear versus cytoplasmic mRNA noise (from smFISH) for all promoters (LTR, UBC, EF-1α, and SV40) shows that noise is primarily amplified from nucleus to cytoplasm. Data points are mean of two biological replicates, and error bars represent SEM.

**Figure 3:**
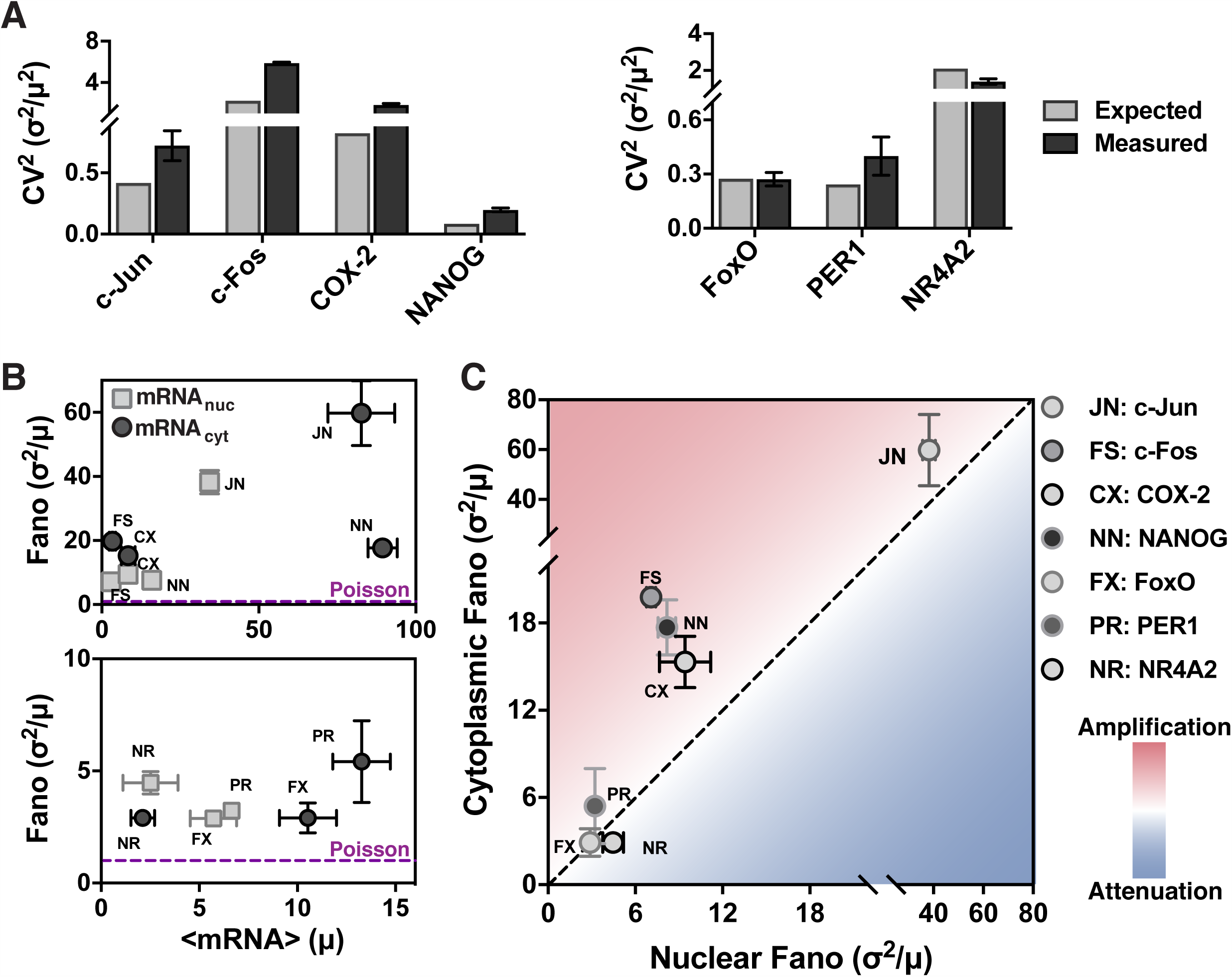
Endogenous genes exhibit amplification of mRNA noise from nucleus to cytoplasm. (**A-B**) smFISH analysis, post extrinsic-noise filtering, for Per1, NR4A2, FoxO, c-Jun, c-Fos, and COX-2 mRNAs in human embryonic kidney cells (293) and for NANOG mRNA in mouse embryonic stem cells. Data points are biological replicates, and error bars represent SEM. (**A**) Expected versus measured cytoplasmic CV^2^ of expressed mRNAs. (**B**) Mean mRNA expression (μ) versus noise (σ^2^/μ) for both nuclear (squares) and cytoplasmic (circles) mRNAs. (**C**) Comparison of nuclear versus cytoplasmic noise, (from smFISH) for Per1, NR4A2, FoxO, c-Jun, c-Fos, COX-2, and NANOG shows that cytoplasmic mRNA noise is primarily amplified.

For the reporter constructs, lentiviral vectors were used to semi-randomly integrate the reporters into the genome and isoclonal populations were generated from individual transduced cells such that each different isoclone carried a single promoter integrated at a unique genomic locus. Importantly, this approach controls for the effect of a specific genomic locus on, for example, noise levels (Becskei et al., 2005) or localization of nuclear transport machinery (Casolari et al., 2004) by allowing the same promoter to be analyzed at multiple genomic loci. The reporter constructs (Figure 2A) used a range of both human and viral promoters, including: the human ubiquitin C (UBC) promoter, which drives an essential cellular housekeeping gene and results in abundant protein expression across integration sites and cell types (Kim et al., 1990); the human elongation factor 1α (EF-1α) promoter, a stronger constitutive promoter also expressing an essential cellular housekeeping gene; the HIV-1 long terminal repeat (LTR) promoter, an inducible and exceptionally bursty viral promoter; and, the Simian virus 40 (SV40) promoter, a viral promoter that is far less noisy than the LTR promoter (Dar et al., 2012; Gilbert et al., 2013).

Isoclonal populations were imaged using three-dimensional (3D) confocal microscopy (Figure 2B and S2A–B), and individual mRNA molecules were quantified with a series of extrinsic-noise filtering steps that eliminate contributions to the Fano factor arising from external stimuli (Raj et al., 2006). Consistent with previous observations (Padovan-Merhar et al., 2015), analysis of correlation strength between cellular volume, shape, and DNA-stain intensity with mRNA count indicated that cell size was the strongest measure of extrinsic noise (i.e., mRNA copy number scales most tightly with cell size) (Figures S2C–G). Consequently, our analysis focused primarily on size-dependent extrinsic-noise filtering (Figures S2D-G), as done in similar genome-wide analyses in yeast (Newman et al., 2006). Nuclear and cytoplasmic mRNA counts were measured through 3D image analysis and DAPI staining of nuclear DNA. Frequency distributions (Figure 2C) were obtained for each isoclonal population of cells (minimum of ~100 cells, as lower cell counts significantly increased the calculated Fano factors by increasing the effect of outliers) (Figure S2E).

As predicted by the simulations (Figure 1) and analytic arguments (Eq. [1]), smFISH mRNA quantification largely show amplification of mRNA noise in the cytoplasm relative to the noise in the nucleus; in virtually all cases, the CV^2^ of cytoplasmic mRNA is significantly higher than expected from Poisson scaling (Figure 2D and S3A), with all data falling far above the minimally stochastic Poisson noise (Fano factor = 1) for all promoters (Figures 2E). The SV40 promoter was the only promoter with comparable mRNA noise in the nucleus and cytoplasm, although it still generated mRNA noise far from Poissonian in both the nucleus and cytoplasm. Directly comparing cytoplasmic versus nuclear noise for all four promoters (16 genomic loci), shows that, in most cases, cytoplasmic noise was significantly amplified relative to nuclear noise (Figures 2F), with the data falling within the parameter space that most genes were predicted to fall (Figure S3B). This generalized amplification of mRNA noise in the cytoplasm occurs despite very different mRNA mean and noise levels, consistent with transcriptional burst size and frequency affected by genomic location (Dar et al., 2012).

To further validate these results, smRNA FISH was performed on seven endogenous genes in adherent cell lines (mouse embryonic stem cells and human embryonic kidney cells). These measurements encompass five signal-responsive genes (three immediate-early response genes c-Jun, c-Fos, and NR4A2; the forkhead transcription factor, FoxO1; and, a late-response gene COX-2), a circadian clock gene (PER1), and a gene constitutively expressed in the pluripotent state (NANOG). Consistent with data from non-adherent cell lines (Figure 2), the majority of these genes exhibit cytoplasmic mRNA noise that is amplified relative to nuclear noise (i.e., CV^2^ larger than expected from Poisson scaling) and is far above the minimally stochastic Poisson limit (Figure 3A-B and S3C). While NR4A2, COX-2 and c-Fos have a similar number of mRNAs in the nucleus and cytoplasm, NANOG, PER1, FoxO and c-Jun have higher cytoplasmic than nuclear means (Figure 3B). Five of the genes show higher cytoplasmic than nuclear noise and fall in the amplification regime, FoxO falls in the unchanged regime, and NR4A2 is the only gene which shows slight attenuation of cytoplasmic mRNA noise compared to nuclear mRNA noise (Figure 3C). Thus, in agreement with theoretical predictions (Figure 1 and Eq. [1]), experimental observations show that the majority of genes exhibit amplification of mRNA noise in the cytoplasm compared to the nucleus, especially when the mean cytoplasmic abundance is greater than its mean nuclear abundance.

### As predicted, cytoplasmic mRNA and protein noise are largely insensitive to changes in nuclear export

To test model predictions of the effects of nuclear export on cytoplasmic noise, we computationally and experimentally perturbed nuclear export rates. Numerical simulations and analytical arguments (Eq. [1]) predicted that slowed nuclear export should only impact nuclear noise, without affecting cytoplasmic noise (Figure 4A, Figure S4A), because most genes fall in a regime in which nuclear export is much faster than cytoplasmic mRNA degradation (*k_exp_* >> *k_deg_*). An important assumption of the model is that nuclear-export rate is not operating in the saturated regime, which could lead to nuclear pileup of mRNA, manifesting as reduced net export, and alter these predictions (Xiong et al., 2010). To verify that nuclear export is not saturated, transcriptional center (TC) intensity and frequency were measured by smFISH, and then used together with nuclear mRNA means to quantify the export rate (see Methods: Rate calculations) after transcriptional activation with tumor necrosis factor (TNF) (Duh et al., 1989) for 24 hours. We did not observe altered nuclear-export kinetics compared to the untreated control, indicating that export is operating far from saturation (Figure S4F). Overall, simulations predict that export rates can be the cause of altered nuclear-to-cytoplasmic noise ratio (Figure 4A); however, the export rate does not impact cytoplasmic mRNA noise and, consequently, is predicted to have no impact on protein noise.

**Figure 4:**
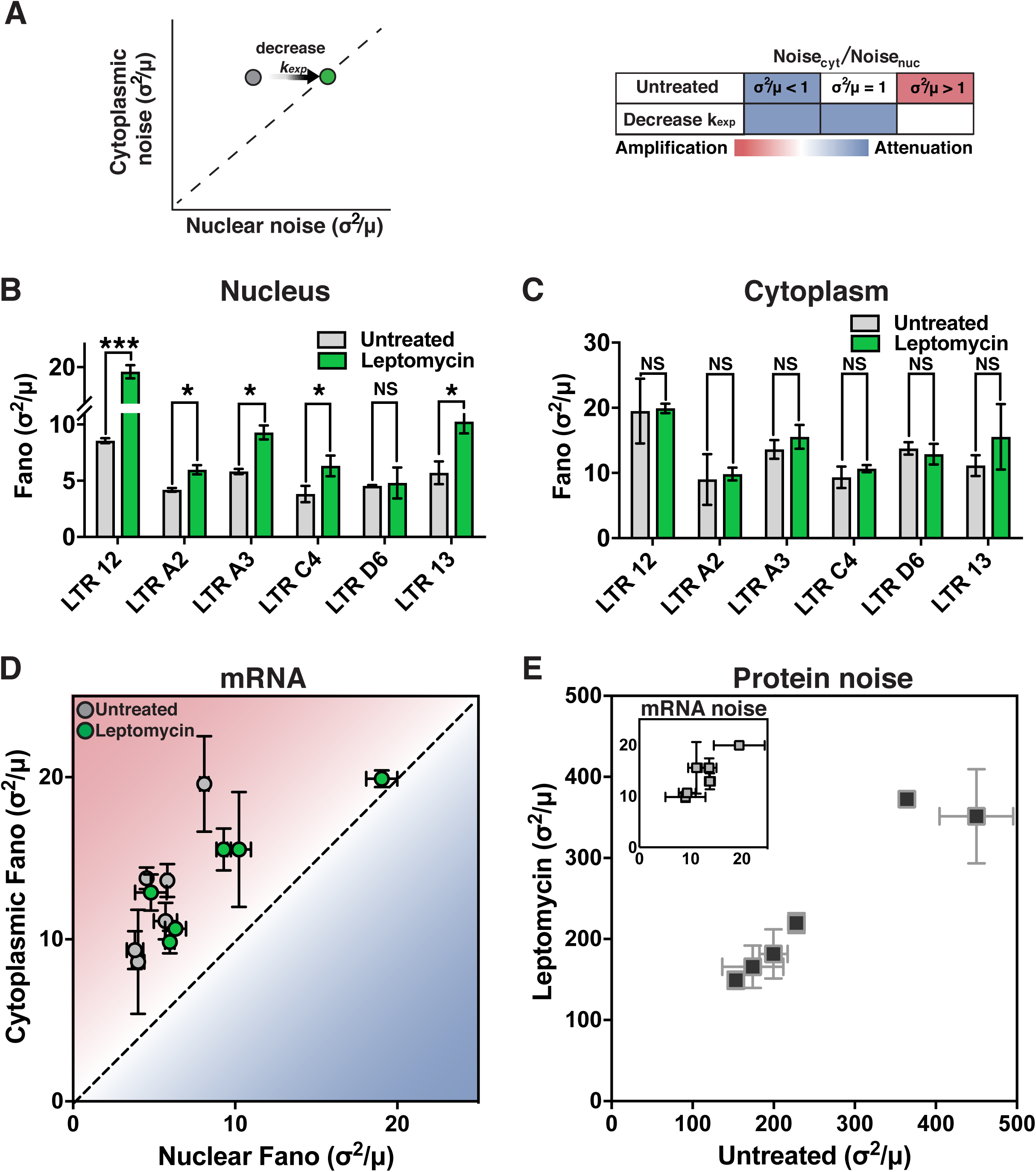
Slowed nuclear export can cause apparent attenuation of nuclear-vs-cytoplasmic RNA noise by amplifying nuclear RNA noise and not decreasing cytoplasmic RNA noise. (**A**) Simulations predict that slowing the nuclear export rate shifts the nuclear-to-cytoplasmic noise ratio by affecting nuclear noise. (**B–C**) smFISH analysis of HIV LTR–expressed mRNA in isoclonal cells treated with the nuclear-export inhibitor leptomycin B (green). Nuclear mean and noise increase whereas cytoplasmic mean or noise remain unchanged (grey). (**D**) Comparison of nuclear versus cytoplasmic mRNA noise by smFISH analysis before and after leptomycin B treatment. All isoclonal populations remain in the amplification regime. (**E**) Protein (d_2_GFP) noise of the same isoclones measured by flow cytometry before and after 5 hours of leptomycin B treatment. As predicted, no change in noise is observed. Inset: cytoplasmic mRNA noise from (C) for comparison. All data points represent means of two biological replicates, and error bars represent SEM.

To experimentally perturb nuclear-export rates, we took advantage of the fact that HIV and lentiviral reporter constructs (e.g., the LTR-GFP construct, Figure 2A) utilize the cellular chromosome region maintenance 1 (CRM1) pathway for nuclear export (Felber et al., 1989; Malim et al., 1988; Ossareh-Nazari et al., 1997) even in the absence of Rev (Urcuqui-Inchima et al., 2011). Cells were treated with the small-molecule inhibitor of CRM1 mediated nuclear export, leptomycin B (Watanabe et al., 1999), and imaged by smRNA FISH. Both dose and duration of leptomycin B were titrated to determine the maximum tolerable concentration (Figures S4C and S4D) (i.e., 0.6 ng/mL for 2.5 hours gave minimal cytotoxicity while still giving significantly increased mean nuclear mRNA). As above, extrinsic-noise filtering (i.e., cell size and DNA content) was employed, which further controls for cytotoxic effects, because dying and dead cells tend to be smaller. To validate that the nuclear-export rate was specifically decreased, without affecting other rates (Figures S4E), we measured TC intensity and frequency by smFISH, and then used mRNA distributions to calculate rates for each individual biochemical step in mRNA biogenesis, as previously done (Bahar Halpern et al., 2015a; Munsky et al., 2012). To validate the decreased export rate, we confirmed the calculated export and cytoplasmic degradation rates (3.78 ± 0.63 and 0.66 ± 0.14, respectively) by smRNA FISH at 15-minute time intervals after treatment with an orthogonal transcriptional inhibitor (Figures S4F), Actinomycin D (Bensaude, 2011), and found an export rate of 4.4 ± 1.5 hr^-1^ and degradation rate of 1.0 ± 0.5 hr^-1^.

As predicted, the data shows that both the mean and noise of nuclear mRNA increased significantly (paired t test: p=0.012 and p=0.007) when nuclear export is diminished approximately 2-fold (Figure 4B). In contrast, neither mean nor noise of cytoplasmic mRNA change significantly when nuclear export is diminished (paired t test: p = 0.374 and p = 0.06, respectively) (Figures 4C and 4D). Protein noise did not change significantly (paired t test: p = 0.162) (Figure 4E) despite use of a short-lived GFP reporter that is particularly sensitive to changes in noise (Dar et al., 2012). This result is expected given the lack of change in cytoplasmic mRNA noise. To test that these results were not caused by an off-target effect of leptomycin B (i.e. observed effects were specific to inhibition of CRM1 export pathway), we treated the SV40 and UBC promoters with leptomycin B. Transcripts expressed from either promoter lack an RRE (Figure 2A) and are presumably not exported via the CRM1 dependent pathway. Therefore, mRNA expressed from either SV40 or UBC should be insensitive to leptomycin B treatment. As expected, we found no significant difference in nuclear mRNA distributions (KS test p > 0.1 compared to p <0.0001 for LTR isoclone A3, Figure S4G).

In extreme cases when export rates fall below the mRNA-degradation rates—as occurs for a small fraction of genes (Bahar Halpern et al., 2015a)—the situation is slightly different. In this regime, cytoplasmic noise can be affected by decreases in export rate (Figure S4A). However, even for a conservatively low protein half-life of two hours, simulations show that this decrease in cytoplasmic mRNA noise cannot propagate to protein noise (Figure S4B). Consequently, for physiologically relevant parameters, even in the extreme cases, nuclear export is predicted to cause noise amplification rather than attenuation.

### Cytoplasmic mRNA noise is further amplified by super-Poissonian mRNA decay and translation processes caused by mRNA switching between alternate states

Based on previous reports (Battich et al., 2015), we next explored whether there might be noise-attenuation processes concealed within the data. Briefly, we used a common model-validation approach (Munsky et al., 2012) to determine whether cytoplasmic mRNA distributions could be predicted from measured nuclear mRNA distributions using the existing model parameter estimates (Figure S4E). If nuclear mRNA distributions predicted broader cytoplasmic mRNA noise distributions than experimentally measured, it could indicate hidden noise-attenuation processes. Importantly, the goal of this analysis is distinct from the analysis above (Figure 2), which shows that cytoplasmic noise is higher than predicted from the cytoplasmic mean level: instead, the goal of this analysis was to test if measured cytoplasmic noise levels are different than predicted from nuclear parameters (i.e., transcriptional burst frequency, transcriptional burst size, mean, and noise).

Strikingly, the data show precisely the opposite of attenuation: the experimentally measured cytoplasmic RNA distributions are significantly broader than predicted from nuclear distributions [KS test: p = 0.0002 (Figure 5A, right panel, grey bar chart versus green dashed line)]. To fit both nuclear and cytoplasmic mRNA distributions, we then analyzed a series of models of increasing complexity in order to arrive at a best-fit model of lowest complexity (Figure S5). We examined eight models consisting of: (Models i-iii) single to multiple mRNA states followed by first order mRNA degradation; (Models iv-v) single to multiple mRNA states with zero-order mRNA degradation; (Models vi-viii) multiple mRNA states with the rate of mRNA entering the degradation-competent state exhibiting zero-order kinetics, followed by first order mRNA degradation. While nuclear RNA distributions could be fit by all models examined, overall, the key physiological process required to fit the amplified RNA noise in the cytoplasmic was that mRNA degradation not be a simple Poisson process (i.e., exponential waiting times), but rather at least a three-state process where degradation is biphasic—in general, multi-state processes are super-Poissonian processes generating super-Poissonian distributions (Singh et al., 2010). Interestingly, we found that the reported biphasic mRNA decay (Yamashita et al., 2005), required that the rate of mRNA entry into the degradation-competent state have zero-order kinetics (Figure S5vi-viii), indicating a reaction rate that is independent of the concentration of the mRNA species involved (Figure S5, blue arrow). In fact, the resulting best-fit model [Model viii; KS test: p = 0.44 (Figure 5A, right panel, grey bar chart versus purple full line)] incorporates two previously documented phenomena: (i) biphasic mRNA degradation in mammalian cells (Yamashita et al., 2005)—where two independent co-translational deadenylation steps (state 1 and state 2) are followed by 5′ decapping (state 3) and mRNA degradation; and (ii) the well-documented inverse relationship between translational rates and rates of mRNA degradation (LaGrandeur and Parker, 1999; Parker, 2012), which posits that translational machinery protects mRNA from degradation or, vice versa, that the presence of mRNA degradation machinery inhibits translation.

**Figure 5:**
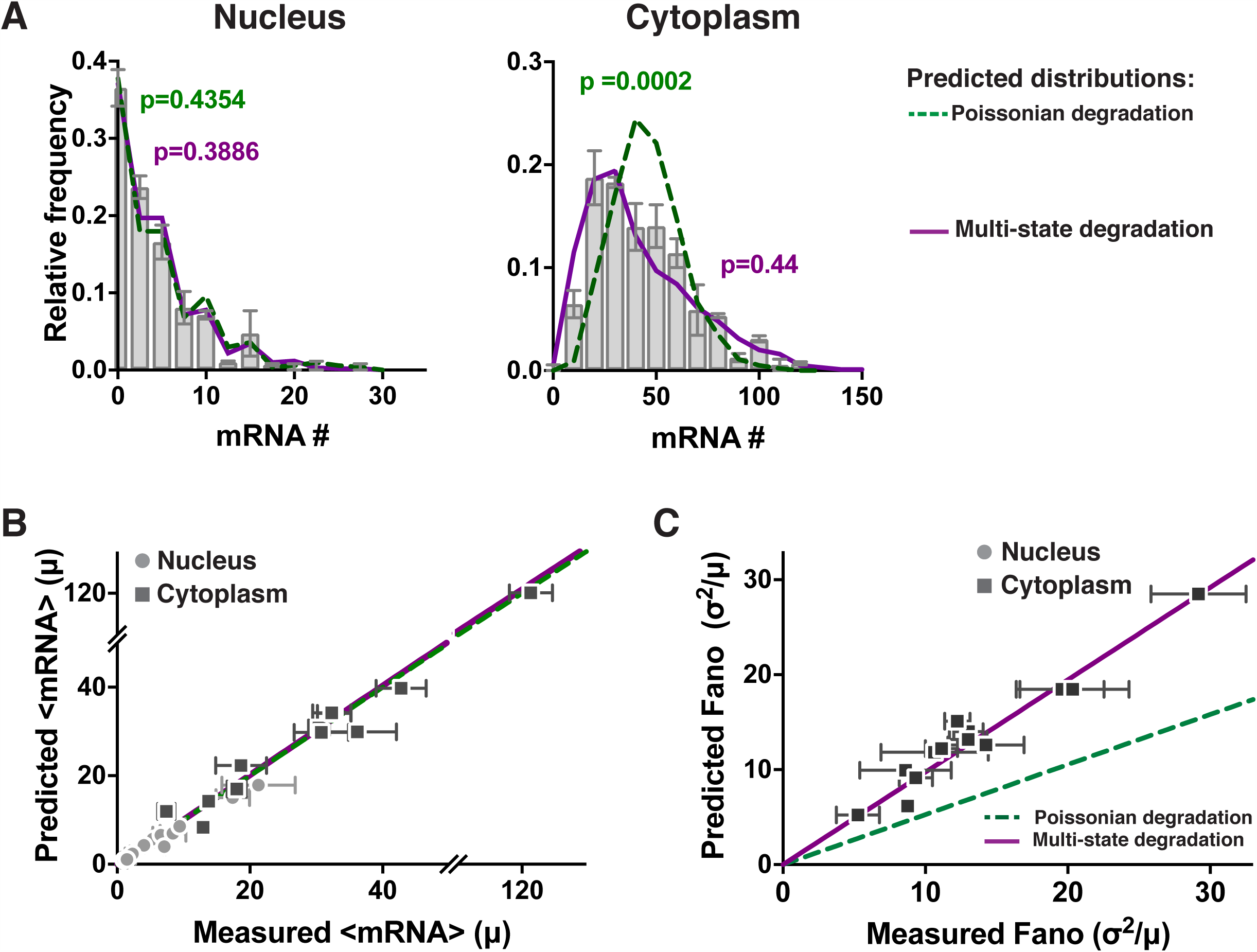
Cytoplasmic mRNA noise is further amplified by multi-state mRNA decay. (**A**) Representative smFISH probability distributions of nuclear and cytoplasmic HIV LTR–expressed mRNA in an isoclonal population of human T lymphocytes. Both the single-state Poissonian degradation model (dashed green line) and multi-state degradation model (solid purple line) fit the experimental probability distribution (bar graph) of nuclear mRNA levels (left column), but the mRNA distribution in the cytoplasm (middle column) is significantly wider than predicted from Poissonian degradation (dashed green line) and fits a multi-state super-Poissonian degradation model (solid purple line). Schematics of each degradation model are shown on the right. P values are from KS test to the experimental data. (**B**) Both the super-Poissonian and Poissonian degradation models accurately predict nuclear (circles) and cytoplasmic (squares) mean mRNA levels – paired t-test p = 0.1623 and 0.3737 respectively. (**C**) The Poissonian degradation model (dashed line) does not accurately predict the cytoplasmic mRNA noise – paired t-test p=0.0002. Only the super-Poissonian degradation model (solid line) accurately predicts both nuclear (circles–inset) and cytoplasmic (squares) mRNA noise – paired t-test p=0.1411 and 0.1623 respectively. Inset: both models accurately predict nuclear mRNA noise. (**B–C**) All data points are mean of two biological replicates, and error bars represent SEM. Lines are linear regressions.

To further validate the multi-state degradation (super-Poissonian) model, we analyzed the panel of clones from above (Figure 2). We first double-checked that the measured TC frequency and size could accurately predict mRNA distributions in the nucleus and, consistent with the computational predictions, measured TC size and frequency indeed accurately predicts nuclear mRNA distributions using either a Poisson or non-Poisson model for all clones (Figures 5A, left panel, and 5B-C circles). However, consistent with the results above (Figure S5), for all clones examined, cytoplasmic mRNA distributions had substantially higher noise than predicted by Poisson degradation and these super-Poissonian distributions could be fit by the multi-state mRNA degradation model (Figures 5A, right panel, and 5C squares). Overall, these data confirm a further amplification of cytoplasmic mRNA noise relative to the nucleus, consistent with multi-state (super-Poissonian) mRNA degradation.

To experimentally test the multi-state translation-degradation model, we next analyzed the effects of two small-molecule inhibitors (cyclohexamide and lactimidomycin) that block mRNA translation through alternate mechanisms of action (Figure S6A-C) and that numerical simulations predicted would have inverse effects on mRNA half-life in the translation-degradation model (Figure S6D, dashed lines). Specifically, cyclohexamide (CHX) inhibits the elongation of ribosomes, causing ribosomes to accumulate on the mRNA (Lee et al., 2012), whereas lactimidomycin (LTM) inhibits the final step of translational initiation (Lee et al., 2012). If mRNAs undergo multi-state degradation-translation with ribosomes protecting mRNAs from degradation, the model predicts that CHX would prevent transcripts from entering the degradation-competent state, resulting in a lower *k_on_deg_* and longer mRNA half-life (predictions shown in Figure S6D, left – dashed blue line). In contrast, in the presence of LTM, the mRNA is free of ribosomes—except for the initiating ribosome which is frozen in place—and more susceptible to exosomal decay (Garneau et al., 2007). Consequently, the model predicts that LTM should push transcripts into the degradation-competent state, resulting in a higher *k_on_deg_* and a decrease in mean cytoplasmic mRNA per cell (Figure S6D, right – dashed red line). Strikingly, despite both CHX and LTM inhibiting protein translation to the same extent (Figure S6A), CHX causes an accumulation of cytoplasmic mRNA over time while LTM treatment shows a decrease in cytoplasmic mRNA (Figure S6D), as predicted. We did observe LTM inducing an initial 1-hr transient increase in cytoplasmic mRNA preceding the decrease, and, as previously reported, this transient increase could be due to the cell globally decreasing degradation rates as a response to stress (Horvathova et al., 2017). Importantly, these changes in cytoplasmic mRNA levels are not due to changes in transcription rate, since the nuclear mean (Figure S6D, black data points) and respective distributions (Figure S6B) show no significant differences. Overall, a multi-state mRNA translation-degradation model appears to be the most parsimonious with the cytoplasmic RNA data.

### Multi-state mRNA translation-degradation amplifies protein noise, accounting for up to 74% of intrinsic cell-to-cell variability in protein levels

To examine how mRNA noise propagates to protein levels, we combined quantitative protein imaging with cytoplasmic smRNA FISH. GFP levels were quantified in individual isoclonal LTR-GFP reporter cells by confocal microscopy (Figure 6A-B, grey circles and bars), and molecular concentrations were calculated by calibration against purified, soluble GFP standards (Figure S7A). Using the measured mean GFP level and half-life (Dar et al., 2012), in combination with previously established parameters (Figures S4E), simulations were used to examine the models of increasing complexity (Figure S5). Remarkably, as above, the protein distributions can only be fit using the eighth model where both degradation and translation are multi-state (i.e., super-Poissonian) processes (KS test p = 0.46). Notably, this multi-state process is fundamentally different from previously reported translational “bursting” (Thattai and van Oudenaarden, 2001) (Figure S7C, green line), as the zero-order entry to a decay-competent mRNA state generates a significantly higher degree of noise amplification in a translation-competent mRNA species and hence in protein (Figure S7C, purple line). This multi-state degradation and translation model—where translation and degradation are mutually exclusive—appears necessary and sufficient to explain the amplified mRNA noise in the cytoplasm, as well as the measured protein noise for the various promoters and integration sites examined (Figure 6A-B).

**Figure 6:**
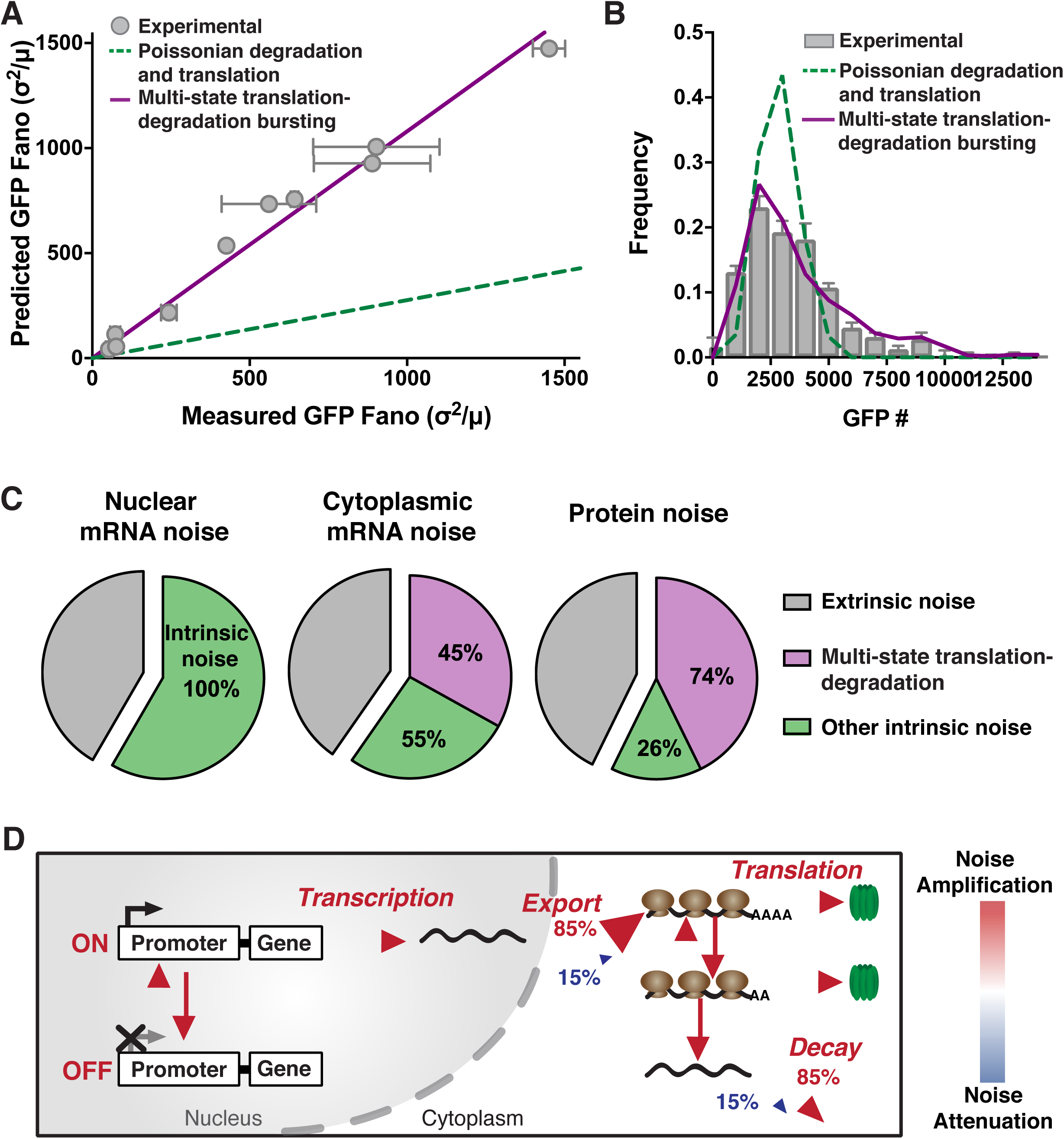
Protein noise is linked to cytoplasmic mRNA noise, indicating an overall model for amplification of transcriptional noise. (**A**) Measured versus predicted noise (σ^2^/μ) of d_2_GFP levels expressed from the HIV LTR promoter in isoclonal population of human T lymphocytes, and from the UBC, and EF-1α promoters (grey squares) in isoclonal populations of human myeloid leukemia cells, determined by microscopy. The single-state Poissonian translation-degradation model (dashed line) poorly predicts the measured d_2_GFP expression noise (grey circles). While the super-Poissonian translation-degradation model (solid line) can accurately predict protein noise. Data points are the mean of two biological replicates, and error bars represent SEM. (**B**) Representative probability distribution of experimental data (bar graph) is significantly wider than the distribution predicted from Poissonian degradation and translation (dashed line), but is fit by a multi-state degradation and translation model (solid line). P values are from KS test to the experimental data. (**C**) Contributions from extrinsic noise (grey), multi-state translation-degradation (purple) and all other intrinsic noise (green) to total nuclear mRNA, cytoplasmic mRNA or protein noise. (**D**) Schematic of cumulative model showing steps that amplify (red) or attenuate (blue) expression noise.

Next, to determine the contribution of multi-state translation-degradation bursting to overall cellular noise, and specifically intrinsic noise, we analyzed flow cytometry data and microscopy measurements against predictions from numerical simulations. First, using an established size-gating approach (Blake et al., 2003; Newman et al., 2006; Singh et al., 2010) that isolates cells synchronized by cell-cycle state, we filtered intrinsic from extrinsic noise (Figure S8A and S2C-G). While this size-gating approach may slightly under-estimate extrinsic factors (Newman et al., 2006) it excludes the majority of extrinsic cellular variability especially for low expressing genes, an expression regime where other methods (e.g., two-color analysis) are technically difficult to employ. Consistent with previous large-scale analysis of low abundance genes in eukaryotes (Bar-Even et al., 2006; Dar et al., 2012; Newman et al., 2006), we find that intrinsic factors appear to account for a large portion of total protein noise in flow cytometry data—the intrinsic contribution ranged from 59–76% for the LTR promoter (Figure S8C and S8E), to 41-61% for more highly expressing promoters (EF-1α, SV40 and UBC, Figure S8B and S8E) and ~37% for constitutively expressed NANOG (Figure S8D and S8E). Then, to determine how translation-degradation bursting quantitatively contributes to this intrinsic noise, we calculated the expected protein noise for all clones (LTR, EF-1α and UBC) under two alternate scenarios: (i) a scenario where noise is generated only from previously characterized sources (i.e., transcriptional bursting, nuclear export, and Poissonian mRNA degradation and translation); or, (ii) the scenario where multi-state translation and degradation is included as a potential noise source. Strikingly, this analysis indicates that, for all cell types and promoters, multi-state translation-degradation bursting accounts for ~74% of intrinsic cell-to-cell variability (Figure 6C) making it the dominant source of intrinsic cellular variability (other processes account for ~26% of intrinsic noise).

## DISCUSSION

How cellular processes amplify or attenuate gene-expression fluctuations (noise) is crucial to designing synthetic gene-regulatory circuits (Hasty et al., 2002) and for efforts to efficiently specify cell fate (Blake et al., 2006; Dar et al., 2014). Here, we analyzed mRNA and protein noise to quantify whether cellular processes amplify or attenuate fluctuations as mRNAs proceed from transcription through translation. Computational results (Figure 1) show that in the majority of physiologically relevant scenarios (approximately 85%), nuclear export of mRNA amplifies mRNA fluctuations generated by transcriptional bursts, and single-molecule RNA counting corroborates this prediction for several viral and mammalian promoters (LTR, UBC, EF-1α, SV40, c-Jun, c-Fos, COX-2, FoxO, Per1, NR4A2 and NANOG) in different mammalian cell types (Figure 2–3). The results also show that cytoplasmic mRNA noise is robust to changes in nuclear export (Figure 4), but can be substantially amplified by super-Poissonian mRNA decay (Figure 5) and translation processes (Figures 6). Cumulatively, the resulting model is capable of predicting protein noise from transcriptional measures and shows that the effects of nuclear export, mRNA degradation, and translation substantially amplify gene expression noise, resulting in cytoplasmic mRNA and protein distributions that are super-Poissonian (i.e., far from minimal Poisson noise). These results, which show that transcriptional noise propagates to the protein level, are consistent with findings that noise exerts purifying (negative) selection pressures on promoter architecture (Fraser et al., 2004; Wolf et al., 2015) and can drive diversifying (positive) selection for bet-hedging phenotypes (Balázsi et al., 2011; Beaumont et al., 2009; Raj and van Oudenaarden, 2008; Rouzine et al., 2015).

These results build on previous findings that translation can proportionally amplify transcriptional fluctuations (Thattai and van Oudenaarden, 2001) since a single RNA typically produces hundreds to thousands of protein molecules (Bar-Even et al., 2006; Blake et al., 2003)—i.e., fluctuations of a single mRNA molecule can generate large fluctuations in protein numbers. However, the multi-state translation-degradation model is fundamentally different from previously reported translational “bursting” (Thattai and van Oudenaarden, 2001), which is insufficient to generate the required noise levels to fit the data. Moreover, the findings herein are consistent with recent studies showing that in ~15% of cases nuclear export is slower than cytoplasmic mRNA degradation, and passive attenuation of mRNA noise can occur (Bahar Halpern et al., 2015). While an accompanying study reported that noise attenuation was more widespread (Battich et al., 2015), the HeLa cells and primary human keratinocytes examined in that study could exhibit significantly slower nuclear export and faster mRNA degradation than the lymphocytes, embryonic murine cells, and kidney cells examined here, which may explain how the same genes (c-Jun, c-Fos, Per1, and FoxO) exhibit different noise properties in these studies. Nevertheless, the results herein demonstrate that any potential attenuation of mRNA noise does not translate to decreased protein noise, due to longer protein half-lives compared to mRNA half-lives, in line with previous predictions (Singh and Bokes, 2012), and possibly due to multi-state degradation and translation (Figure 5–6). Interestingly, a recent study also found higher-than-predicted cytoplasmic noise for transcripts expressed from 12 yeast genes attributed to mRNA processing downstream of transcription and elongation (Choubey et al., 2015). Overall, amplification appears to be the most common form of noise modulation in the absence of specific gene-regulatory circuits.

Most interestingly, these data support a model for cytoplasmic mRNA deadenylation occurring at two distinct rates (Yamashita et al., 2005), with translational initiation and mRNA degradation being inversely proportional and mutually exclusive processes (LaGrandeur and Parker, 1999; Parker, 2012; Pelechano et al., 2015; Schwartz and Parker, 1999). Notably, the model proposed here does not contradict recent data obtained from yeast demonstrating that some mRNA can undergo co-translational mRNA degradation (Pelechano et al., 2015). Since, the model requires translational *initiation* and degradation to be mutually exclusive, elongation could still occur during 5’ to 3’ degradation. Hence, cytoplasmic mRNA is subject to another, at least three-state process, which adds a significant noise-amplification step to gene expression. Several mechanisms could explain this multi-state degradation and translation including the association of non-translating mRNAs into P-bodies or stress granules, as P-bodies and stress granules are enriched with mRNA degradation and translational initiation machinery respectively (Decker and Parker, 2012). However, because our data do not show mRNA aggregates in the cytoplasm, biphasic deadenylation along with mutually exclusive degradation and translation appears to be the more parsimonious model.

While the computational model we employed was admittedly simplified—it considered only two transcriptional states without important processes such as splicing—the smFISH measurements show that amplification of noise occurs primarily via post-transcriptional processes in the cytoplasm (e.g., export, degradation and translation; see Figure 6D). If splicing were included as a rate-limiting step (Hao and Baltimore, 2013), it would add an extra noise source and most likely further amplify noise. Moreover, the results do not depend on the strict two-state random-telegraph transcription model, because noise amplification is primarily post-transcriptional—i.e., the noise amplification result would hold for other transcription models with greater than two transcriptional states (Corrigan et al., 2016; Neuert et al., 2013; Zoller et al., 2015).

It is not clear how widespread biphasic mRNA decay is across the mammalian genome or even across different mammalian cell types, and, notably, mRNAs that exhibit Poisson-like mRNA decay (Horvathova et al., 2017) would not be subject to this additional noise amplification step. From an evolutionary perspective it is conceivable that the ~15% of mRNA species that are subject to passive attenuation of mRNA noise (Bahar Halpern et al., 2015) could also exhibit Poisson-like mRNA decay, allowing for certain genes to have evolved a low-noise gene expression pathway. It also is possible that untranslated mRNAs, such as miRNAs or shRNAs, are not subject to multi-state degradation and accompanying noise amplification, due to the lack of protecting ribosomes, though Dicer and other processing machinery could serve an equivalent protection role. Since miRNAs modulate protein levels via mRNA degradation and translational repression (Bartel, 2004), miRNAs could influence multi-state degradation and translation rates to modulate noise (Garg and Sharp, 2016; Schmiedel et al., 2015).

In summary, the results show that, in the majority of scenarios, transcriptional noise is amplified by nuclear export and is then further amplified by mRNAs switching between translation- and degradation-competent states. Importantly, the results show that this intrinsic cellular process, multi-state translation-degradation, accounts ~74% of the intrinsic noise, providing a foundational basis for noise to have acted as a substrate for promoter selection and as a driving force in cell-fate decisions.

## Author Contributions

M.M.K.H. and L.S.W. conceived and designed the study. M.M.K.H., R.V.D, and L.S.W. designed and performed the experiments. M.M.K.H., M.L.S. and L.S.W. analyzed the data and models. M.M.K.H. and L.S.W. wrote the paper.

## Acknowledgments

We are grateful to Brandon Razooky, Noam Vardi and members of the Weinberger Lab for extensive discussions, as well as Kurt Thorn and DeLaine Larsen (Nikon Imaging Center, UCSF) and the UCSF-Gladstone Center for AIDS Research flow core (funded through P30AI027763 and S10RR028962-01) for invaluable technical expertise. We would like to thank Elizabeth Tanner for the LTR d2GFP isoclones 2, 12, and 13, and Noam Vardi for the LTR d2GFP isoclones A3, C4 and D6, the EF-1α d2GFP isoclones, and the UBC d2GFP isoclones. The K562 cell line, along with the SV40 GFP isoclones, were a kind donation from Jonathan Weissman. M.M.K.H is supported by the Netherlands Organization of Scientific Research (NWO) through a Rubicon fellowship (No. 019.153LW.028). M.L.S. is supported by the Collective Phenomena in Nanophases Research Theme at the Center for Nanophase Materials Sciences, sponsored at Oak Ridge National Laboratory by the Office of Basic Energy Sciences, U.S. Department of Energy. L.S.W. acknowledges support from the Alfred P. Sloan Research Fellowship, NIH awards R01AI109593, P01AI090935, and the NIH Director’s New Innovator Award (OD006677) and Pioneer Award (OD17181) programs.

## METHODS

### Cell lines

Mouse E14 embryonic stem cells (mESCs) were cultured in feeder-free conditions on gelatin-coated, 10cm Corning plates. ESGRO 2i + LIF media (SF016-200) was used for cell culture. Jurkat T Lymphocytes were cultured in RPMI-1640 medium (supplemented with L-glutamine, 10% fetal bovine serum, and 1% penicillin-streptomycin), at 37°C, 5% CO_2_, in humidified conditions at b 0.05 × 10^5^ to 1 × 10^6^ cells/mL. Human immortalized myelogenous leukemia (K652) cells were cultured in RPMI-1640 medium (supplemented with L-glutamine, 10% fetal bovine serum, and 1% penicillin-streptomycin), at 37°C, 5% CO_2_, in humidified conditions at 2 × 10^5^ to 2 × 10^6^ cells/mL. Human embryonic kidney (293FT) cells were cultured in DMEM (supplemented with 10% fetal bovine serum and 1% penicillin-streptomycin) at 37°C, 5% CO_2_, in humidified conditions at 10 to 90% confluency. All cell lines were passaged at least three times prior to smRNA FISH imaging.

### Computational modeling

#### Stochastic simulations

A simplified two-state transcription model incorporating two compartments (nucleus and cytoplasm) was constructed and simulated using the Gillespie algorithm (Gillespie, 1977), with reaction scheme and parameters as defined in Table S1. Stochastic simulations were run in MATLAB. Initial conditions for all species were set to 0, except for Promoter_OFF_ which was set to 1. Simulations were run to time = 25 (arbitrary time units) and 1000 simulations were run for each parameter set. For the final “time-point” of simulations, nuclear and cytoplasmic mean and Fano factor were calculated.

#### Ordinary differential equations for probable parameter regime calculations

Assuming a simplified mathematical model where mRNAs are transcribed at rate α, nuclear and cytoplasmic mRNA means can be approximated by the following equations:

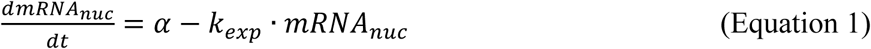

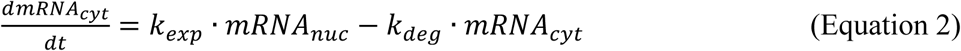

At steady state the mean amount of nuclear and cytoplasmic mRNA is therefore:

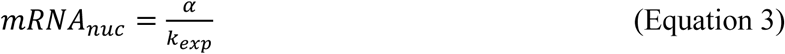

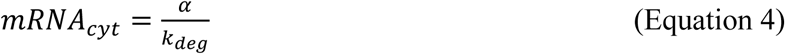

And the ratio of nuclear to cytoplasmic mRNA is:

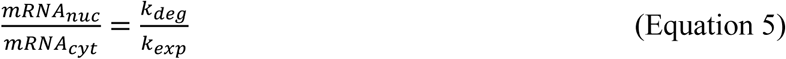

From previously reported genome-wide mRNA counts of nuclear versus cytoplasmic mRNA, the ratio between the degradation and export rate (*k_deg_*/*k_exp_*) per gene can be estimated (Figure S1C) (Bahar Halpern et al., 2015), and the *possible* parameter space can be further limited to a *probable* regime. From the *probable* parameter regime, we could determine how many scenarios resulted in *true* attenuation of mRNA noise (i.e. cytoplasmic noise was attenuated down to minimal Poissonian levels). To remain on the conservative side, a given parameter combination was labeled *true* attenuation, when cytoplasmic noise was lower than nuclear noise, and cytoplasmic Fano factor was < 2. Next, the percentage of scenarios resulting in *true* attenuation was calculated, resulting in only ~2.5%.

#### Analytical derivation

The power spectral density (PSD) of the noise in the Nuc mRNA populations (*S_NUC_*(*f*) is (see Simpson et al. PNAS 2003; Simpson et al. JTB 2004; Cox et al Chaos 2006)

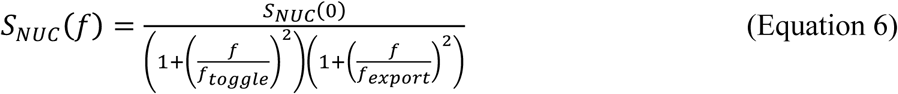

Where

f=frequency in Hz

*S_NUC_*(0) = PSD of NUC mRNA population noise at f=0

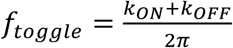 = critical frequency associated with promoter toggling

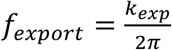 = critical frequency associated with export of mRNA

The variance of the noise in the Nuc mRNA populations 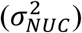 is (Simpson et al. PNAS 2003)

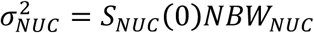

Where

*NBW_NUC_* is known as the noise bandwidth and is a function of the 2 critical frequencies described above (Simpson at al. PNAS 2003).

The Fano factor of the noise in the Nuc mRNA population (FF_NUC_) is

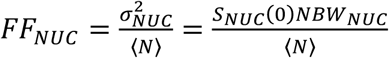

Where

〈*N*〉 is the steady-state value of the Nuc mRNA population.

The PSD of the noise in the Cyto mRNA population (*S_CYTO_*(*f*) is

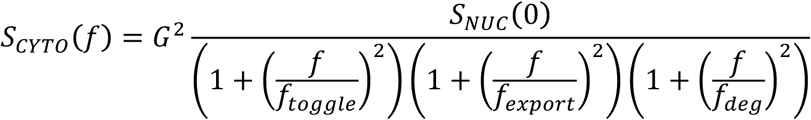

Where

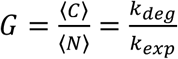 is the “gain” between Cyto and Nuc mRNA populations.

<C> is the Cyto mRNA population

K_deg_is the rate of mRNA degradation in the cyto

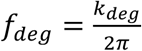 is the critical frequency associated with mRNA degradation in the cyto

The variance 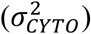 and Fano factors (*FF_CYTO_*) for the Cyto mRNA population are

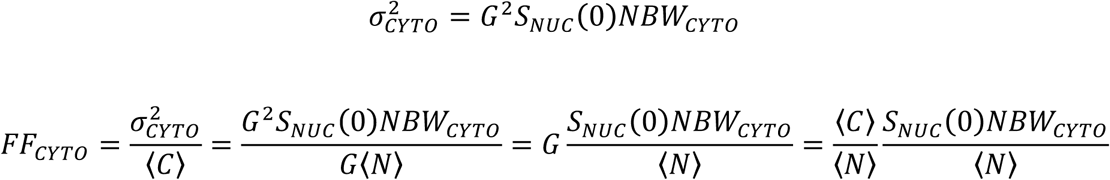

The ratio of the Fano factors is

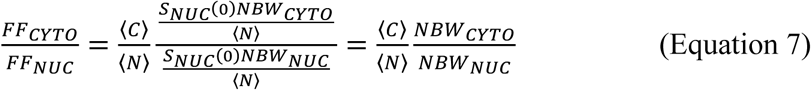

The two noise bandwidths are controlled by the 3 critical frequencies associated with 1. Promoter toggling; 2. mRNA export; and for the cytoplasm 3. mRNA degradation. In both cases, the noise bandwidth is dominated by the lowest of the critical frequencies that are associated with it. As a result,

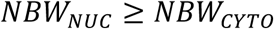

and therefore,

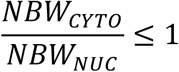

and

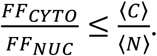

Equation 7 shows that there is a strong tendency for FF_CYTO_> FF_NUC_ for cases where <C> > <N>. Only in the special case where Cyto has a much lower noise bandwidth than Nuc is it possible to have both <C> > <N> and FF_NUC_> FF_CYTO_. These relationships can be made clearer by looking at some limiting cases:

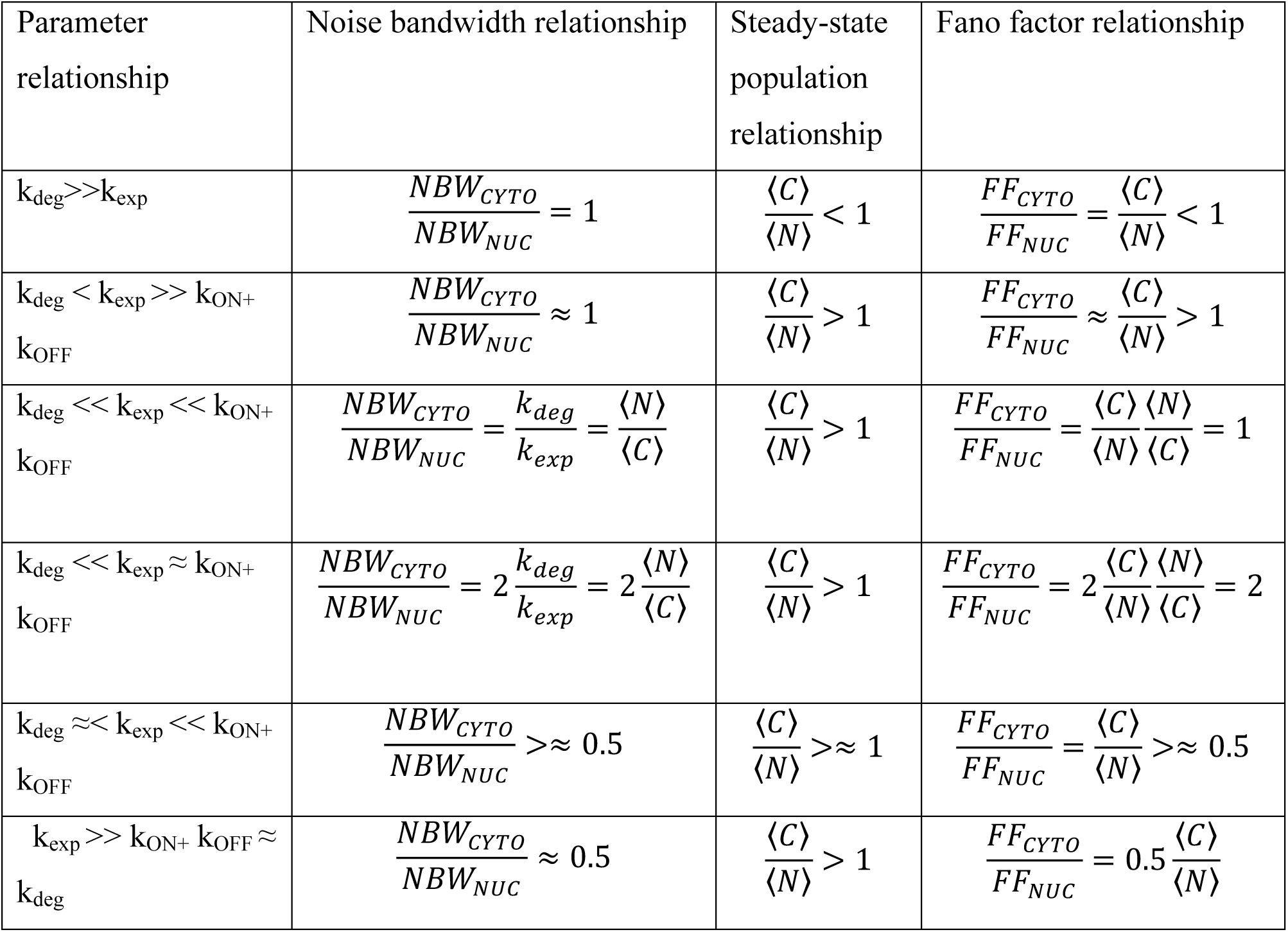

#### Rate calculations

The number of mRNAs at the transcriptional center (*TC_mRNA_*) can be calculated from the transcriptional center intensity (*TC_int_*):

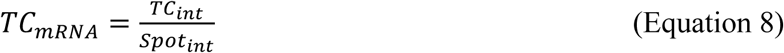

where *Spot_int_* is the median single mRNA intensity. From this the transcription rate (*k_tx_*) was calculated:

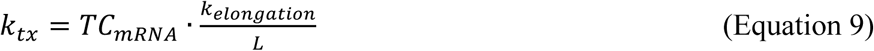

where *k_elongation_* (1.9 kb/min) is the elongation rate of RNA Pol2 (Boireau et al., 2007) and *L* is the length of the gene. The TC frequency is an approximation for the frequency that the respective promoter is on (*f_on_*). Assuming gene expression is at steady-state, then export (*k_exp_*) and degradation (*k_deg_*) rates were calculated from nuclear (*mean_nuc_*) and cytoplasmic (*mean_cyt_*) mean mRNA count respectively (Munsky et al., 2012):

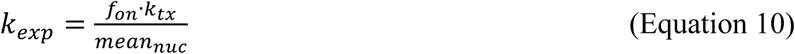

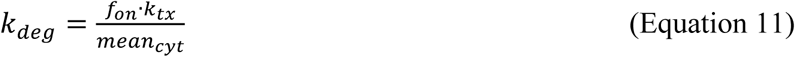

#### Prediction of cytoplasmic noise

From the frequency that the respective promoter is on (*f_on_*):

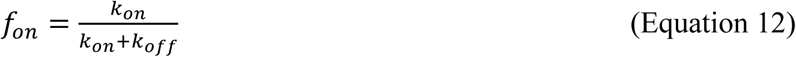

and the nuclear noise (*Fano_nuc_*):

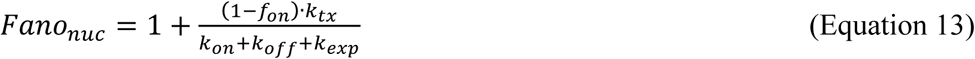

the promoter ON (*k_on_*) and promoter OFF (*k_off_*) rates were calculated (Munsky et al., 2012), given that all other parameters are known. These parameters were then used to predict cytoplasmic noise, using both Poissonian and multi-state cytoplasmic mRNA degradation models. The additional rates involved in multi-state degradation (Table S1) were determined by manual screening of parameter ranges.

### Single molecule RNA FISH

#### Sample preparation

Probes were developed using the designer tool from Stellaris (LGC Biosearch Technologies, Novato, CA) (http://www.singlemoleculefish.com/) to detect GFP, c-Jun, c-Fos, COX-2, FoxO, NR4A2, Per1, c-Fos. Because NANOG had a GFP tag, it could be detected using the GFP probes. Probes were designed using a masking level of 5, and at least 2 base pair spacing between single probes. Each probe set contained 29-48 probes, with each probe being 18-20 nt long and conjugated with TAMRA (see Tables S2–S8 for probe sequences).

Approximately 6×10^5^ isoclonal cells were washed with 2 mL of PBS solution and then immobilized on a Cell-Tak (Corning, Bedford, MA) coated 8-well chambered image dish. Human embryonic kidney (293F) cells were trypsinized with 0.05% Trypsin EDTA (Mediatech, MT 25-052-C1) for 1 minute followed by neutralization with DMEM prior to PBS washing step. If applicable, cells were then treated with leptomycin B (Sigma Aldrich, Darmstadt, Germany). Prior to fixing, mouse E14 embryonic stem cells (mESCs) were cultured on a 35mm MatTek dish (P35G-1.5-14C, MatTek, Ashland, MA). Cells were then fixed with PBS in 3.4% paraformaldehyde for 10 minutes. Fixed cells were washed with PBS and stored in 70% EtOH at 4 °C for a minimum of one hour to permeabilize the cell membranes. Probes were diluted 200-fold and allowed to hybridize at 37 °C overnight. Wash steps and DAPI (Thermo Fisher Scientific, Waltham, MA) staining were performed as described (https://www.biosearchtech.com/support/applications/stellaris-rna-fish).

#### Imaging

To minimize photo bleaching, cells were imaged in a photo-protective buffer containing 50% glycerol (Thermo Fisher Scientific, Waltham, MA), 75 μg/mL glucose oxidase (Sigma Aldrich, Darmstadt, Germany), 520 μg/mL catalase (Sigma Aldrich, Darmstadt, Germany), and 0.5 mg/mL Trolox (Sigma Aldrich, Darmstadt, Germany). Images were taken on a Nikon Ti-E microscope equipped with a W1 Spinning Disk unit, an Andor iXon Ultra DU888 1k x 1k EMCCD camera and a Plan Apo VC 100x/1.4 oil objective in the UCSF Nikon Imaging Center. Approximately 10 xy locations were randomly selected for each isoclonal population. For each xy location, Nyquist sampling was performed by taking ~30, 0.4 um steps along the z-plane. The exposure times for TAMRA (100% laser power), and DAPI (50 % laser power) channels were 500 ms, and 50 ms for single mRNA analysis and 50 ms, and for transcriptional center (TC) analysis. For each z-plane in a 3-D stack images for both single mRNA analysis and TC analysis were taken.

#### Image analysis

mESCs were segmented manually and all other cells were segmented using a short in-house ImageJ script (available upon request), which relied on the auto fluorescence visible in the RFP channel. Spot/TC identification and counting was then performed using in-house MATLAB programs (available upon request). In short, the user enters a cellular size and eccentricity range, DAPI intensity threshold, FISH intensity threshold and TC intensity threshold. The MATLAB program then uses the central z-slice DAPI image to create a general nuclear mask. Together with the cellular mask established in ImageJ this nuclear mask is used to exclude cells outside a given size range, nuclear DAPI intensity range or eccentricity range (see extrinsic noise section below). Cells which contained more than one nuclei were also excluded to eliminate multiple cells which were segmented as one. Notably, these steps automatically exclude unhealthy cells, since the cells tend to shrink and/or DAPI intensity becomes much brighter. Next, the DAPI and FISH images of each individual z-slice are sequentially analyzed. After background subtraction, a Gaussian filter was applied to reduce the amount of local maxima caused by pixel-to-pixel noise. Individual spots and TC’s are then segmented using the predefined thresholds to determine possible spot/TC areas. For each z-slice the local maxima of the segmented spot areas are detected. Local maxima which show up within the possible spot areas of multiple sequential images were only counted as one local maxima. Each DAPI image was used to create a nuclear mask for that specific z-slice, which in turn was used together with the cellular mask to allocate a specific spot to either the nucleus or the cytoplasm of each individual cell. TC’s were defined as such, if their local maxima’s were at least double as bright as the median single mRNA local maxima intensity. The number of mRNAs at the transcriptional center were quantified as the intensity of the transcriptional center divided by the median single mRNA intensity.

#### Extrinsic noise filtering

Extrinsic noise filtering was part of MATLAB program mentioned above, but the respective ranges were determined manually. For the extrinsic noise filtering three parameters were quantified: cellular size; DAPI intensity; and cellular shape (Figure S2C and D). For each parameter the Pearson’s correlation between the total mRNAs per cell and the respective parameter was quantified. The extrinsic noise filtering steps were applied such that the analyzed range of cells was within the respective noise filtering boundaries (Figure S2G). For two extrinsic noise filtering boundaries to be accepted, the Pearson’s correlation must not be statistically significant (i.e. p>0.05, see Figure S2C and F). The final extrinsic noise filtering boundaries were chosen in order to include as many cells as possible while maintaining a Pearson’s correlation p>0.05.

### GFP expression analysis

#### Microscopy

Isoclonal populations were washed with 10 mL PBS solution and then immobilized on Cell-Tak coated 8-well chambered image dish. For the GFP standard curve, soluble eGFP (Cell Biolabs, San Diego, CA) standards (in PBS) of known concentration were imaged under the same conditions as cellular GFP. Both GFP standards, and cellular GFP were imaged on a Nikon Ti-E microscope equipped with a W1 Spinning Disk unit, an Andor iXon Ultra DU888 1k x 1k EMCCD camera and a Plan Apo VC 100x/1.4 oil objective in the UCSF Nikon Imaging Center, exposure time was 200 ms with 20% laser power. Approximately 10 xy locations were randomly selected for each isoclonal population. After background and auto-fluorescence subtraction from cellular GFP images, the cellular GFP concentration was calculated from the GFP standard curve (Figure S6A). Using the measured cellular volume and cellular GFP concentration, the absolute number of GFP molecules per cell was calculated. For a review on molecular counting see (Coffman and Wu, 2014).

#### Flow cytometry

Flow cytometry data was collected on an LSRII cytometer (BD Biosciences) with a 488-nm laser used to detect GFP. The cytometry data was analyzed using FlowJo (http://www.flowjo.com/).

### Quantification and statistical analysis

Statistical analysis was performed by Pearson correlation analysis, Kolmogorov–Smirnov test or paired t test. All data are presented as mean ± SEM or SD. Significance levels were set at *P* < 0.05. For statistical analysis GraphPad^™^ Prism was used.

**Figure S1:**
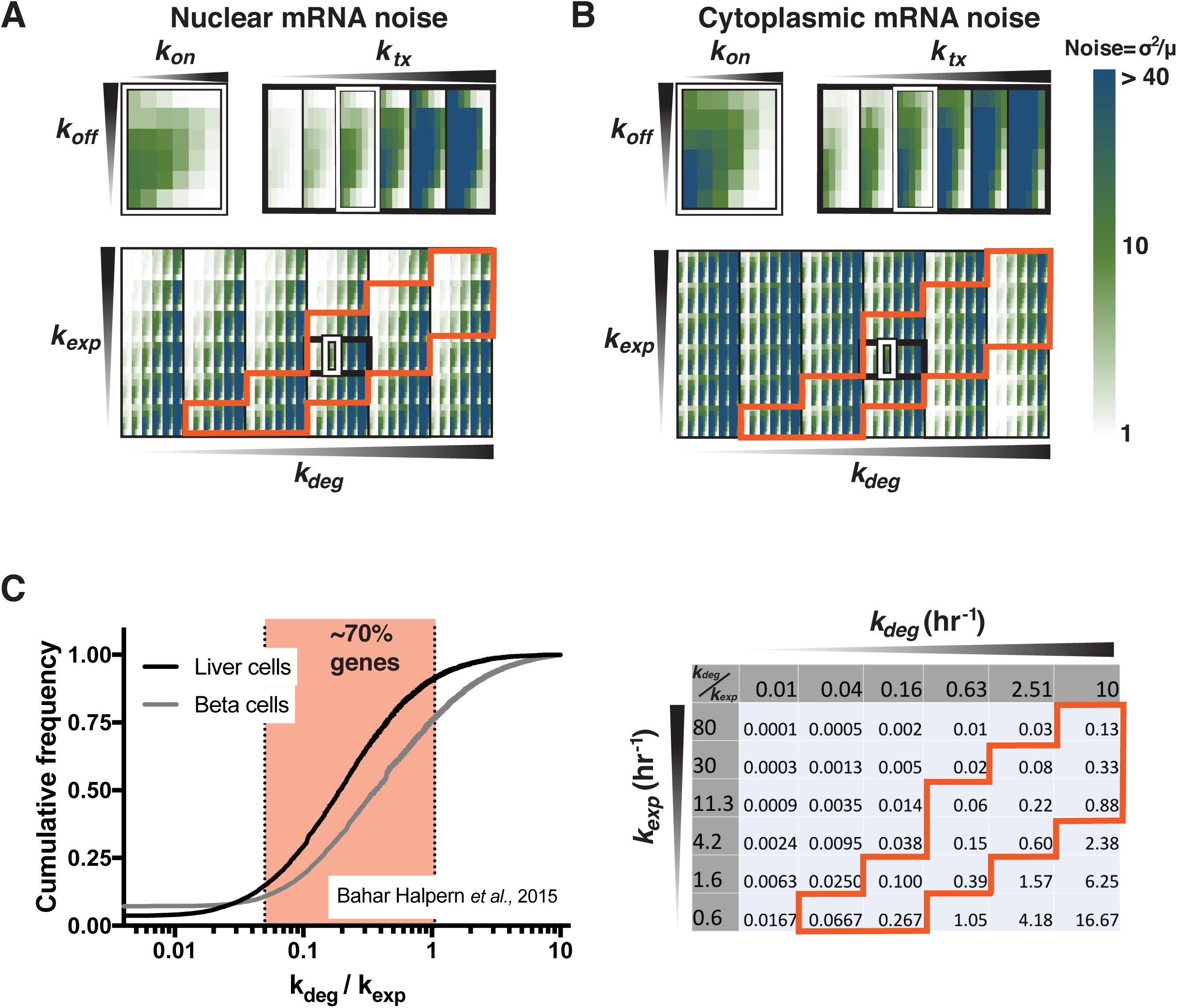
Simulations predicting nuclear and cytoplasmic mRNA noise, related to Figure 1. **(A-B)** Nuclear (A) and Cytoplasmic (B) Fano factor of the physiologically *possible* parameter space, with 1000 stochastic simulations run per parameter combination. The *probable* parameter space is shown in orange. **(C)** Left panel: Cumulative frequency distribution of estimated degradation-to-export ratio (left panel) (Bahar Halpern et al., 2015a). Right panel: *Probable* physiological parameter regime (orange box) was deduced from the cumulative frequency distribution.

**Figure S2:**
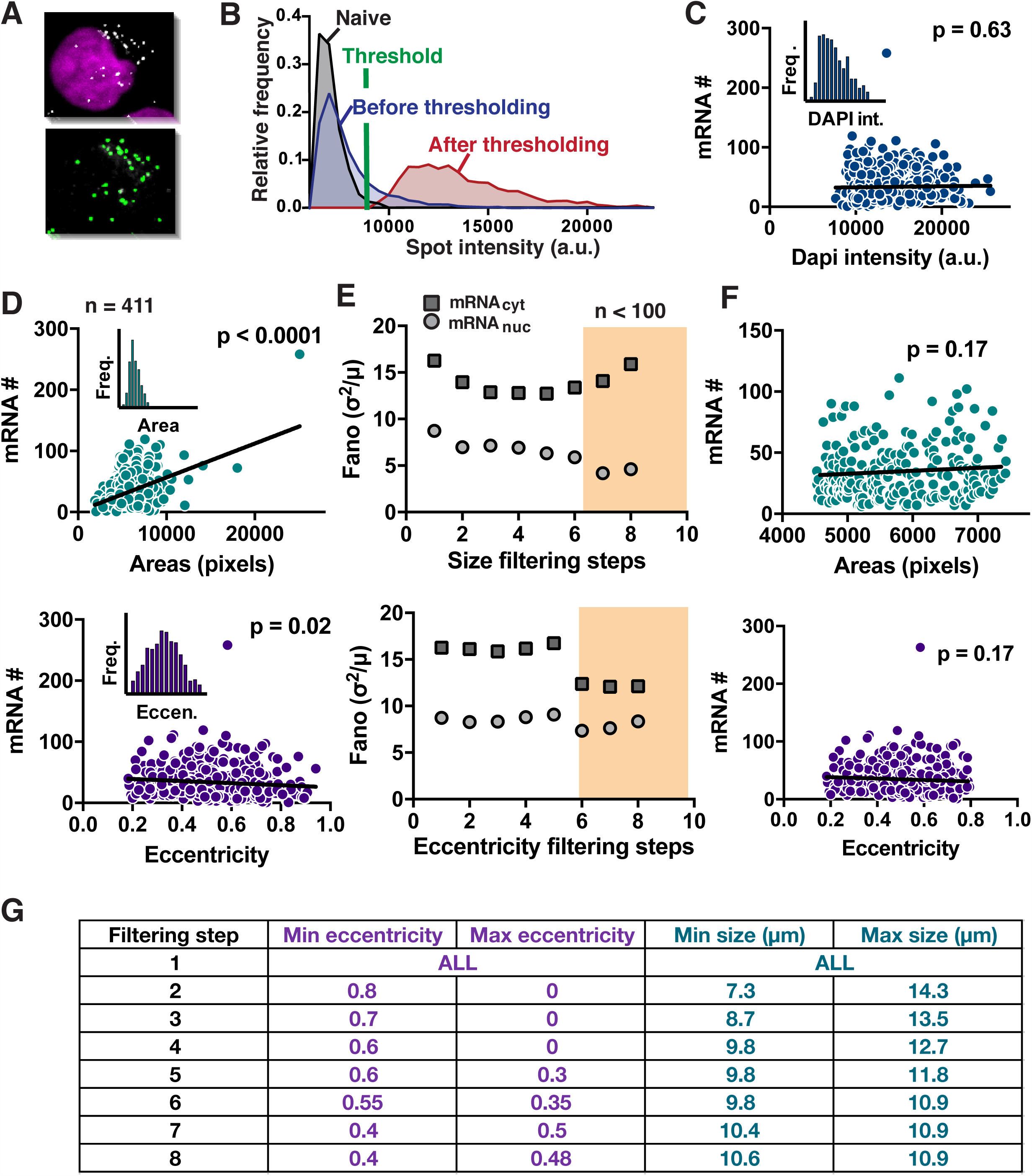
Analysis and of single-molecule mRNA FISH, related to Figure 2. **(A-F)** Isoclonal population of Jurkat cells expressing mRNA from HIV’s long terminal repeat (LTR) promoter. **(A)** Maximum intensity projection of representative smFISH images (top) with mRNA spots marked in green (bottom). **(B)** Probability distribution of mRNA spot intensities of Naïve cells (black), cells expressing the target mRNA before thresholding (blue) and after thresholding (red). **(C-D)** DNA-stain intensity (C), Cellular volume (D, top) and shape (D, bottom), indicating that mRNA copy number scales most tightly with cell size (n=411). Insets: respective probability distributions. **(E)** Extrinsic noise filtering steps using cellular volume (top) and shape (bottom), show that size-dependent extrinsic-noise filtering has the strongest and most consistent effect on mRNA noise. More extrinsic noise steps cause a lower cell count and thus increased the effect of outliers, which can cause higher mRNA noise (shaded area marks < 100 cells). **(F)** The noise filtering steps used for volume (size filtering step #4) and shape (eccentricity step #2), resulted in an insignificant Pearson’s correlation. **(H)** The constraints used for each extrinsic noise filtering step.

**Figure S3:**
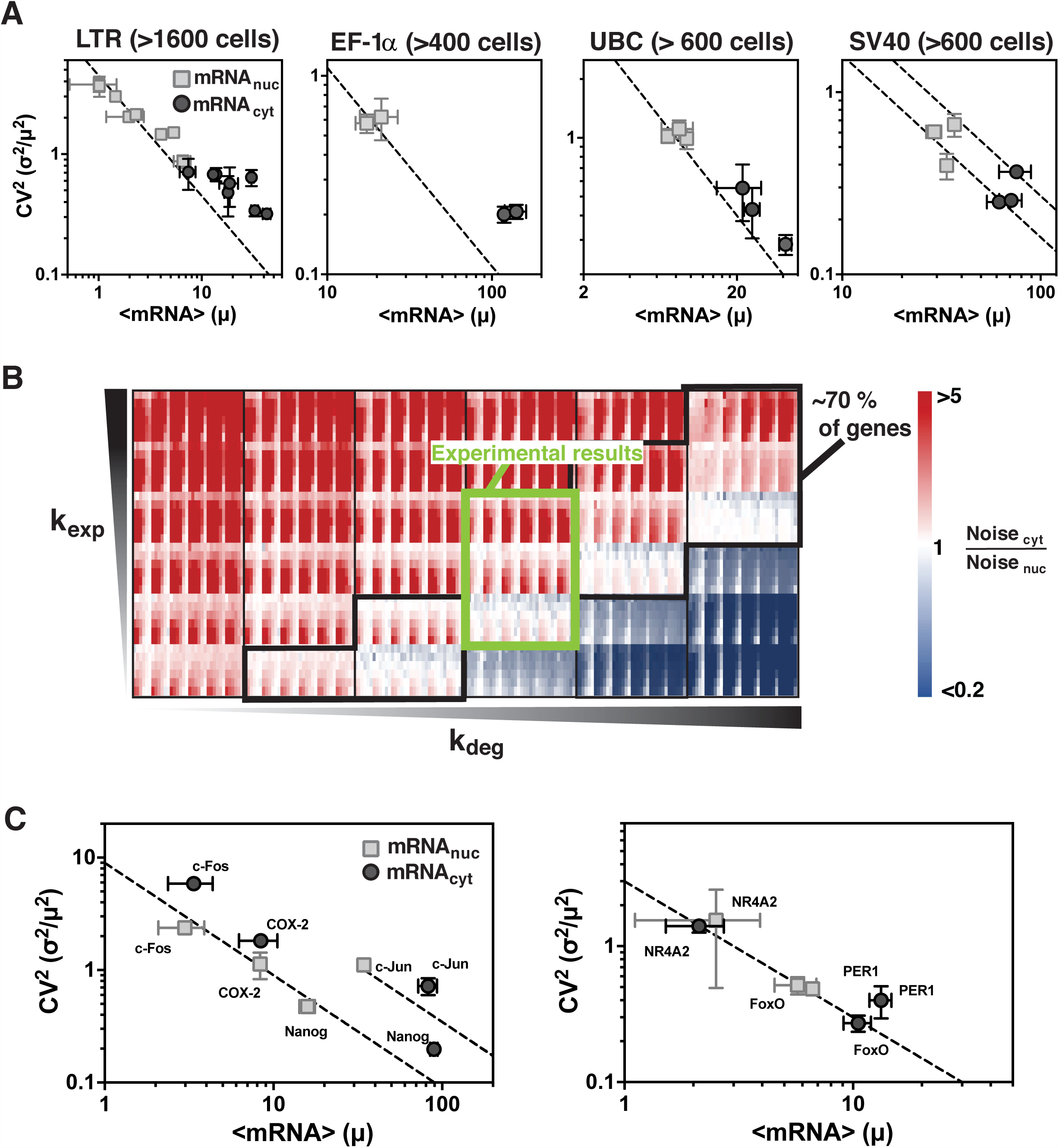
Experimental data falls in the predicted *probable* parameter regime, related to Figure 2. **(A)** Mean mRNA expression (μ) versus CV^2^ (σ^2^/μ^2^) for both nuclear (squares) and cytoplasmic (circles) mRNAs expressed from the HIV-1 LTR promoter in isoclonal populations of human T lymphocytes (Jurkat) and from the UBC, SV40 and EF-1α promoters in isoclonal populations of human myeloid leukemia cells (K562). Data points are biological replicates, and error bars represent SEM. The expected cytoplasmic noise due to Poisson scaling is shown by the dashed line. **(B)** The export and degradation rates of the experimental data (green box) fall within the most *probable* genome-wide parameter regime (black box). **(C)** Mean mRNA expression versus CV^2^ for both nuclear (squares) and cytoplasmic (circles) mRNAs for Per1, NR4A2, FoxO, c-Jun, c-Fos, and COX-2 mRNAs in human embryonic kidney cells (293) and for NANOG mRNA in mouse embryonic stem cells (mESCs). Data points are biological replicates, and error bars represent SEM. The expected cytoplasmic noise due to Poisson scaling is shown by the dashed line.

**Figure S4.**
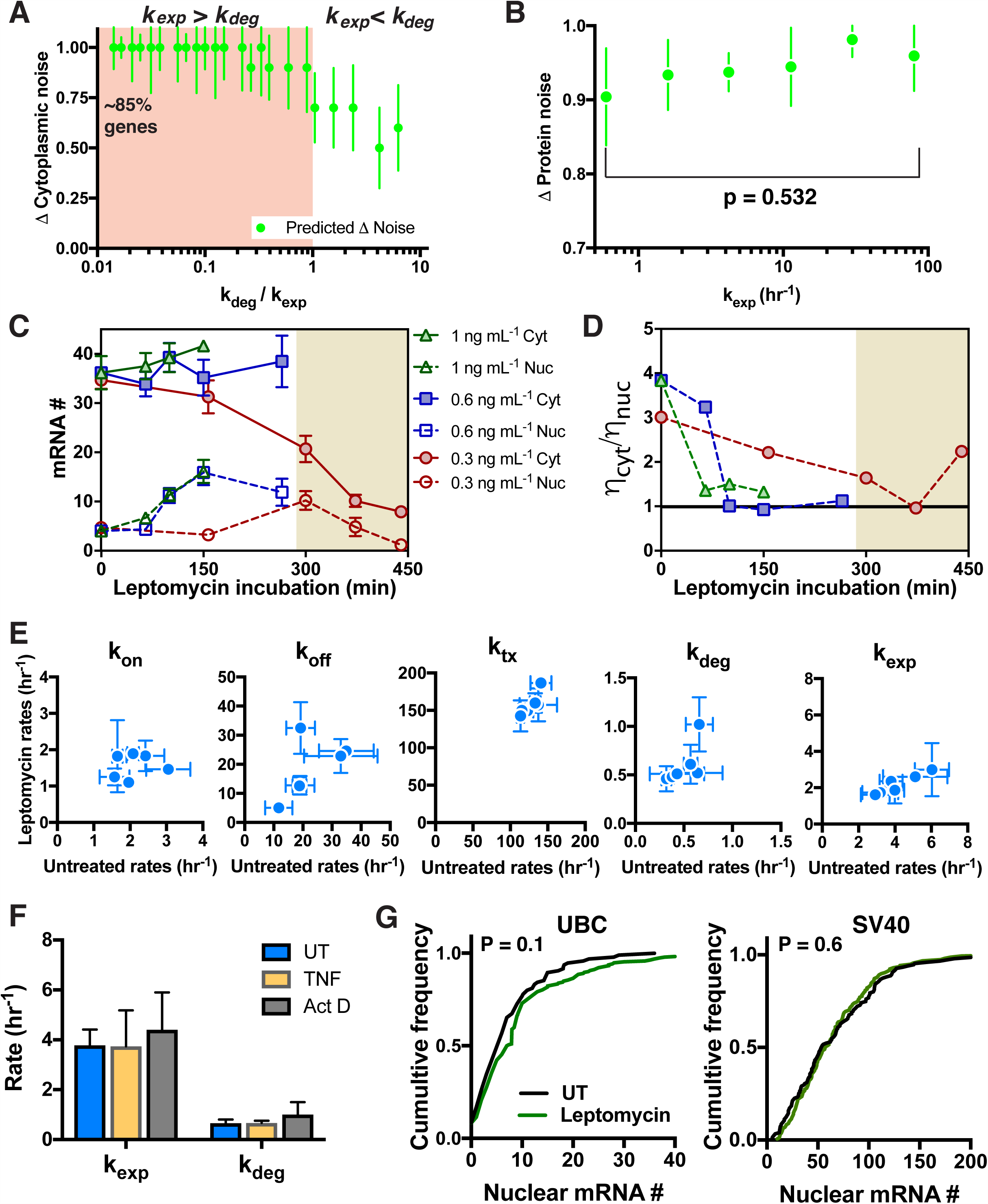
Slowed nuclear export amplifies nuclear RNA noise, related to Figure 4. **(A)** Predicted change (normalized to max) in cytoplasmic mRNA noise when altering only *k_exp_*. Simulations predict that when *k_exp_* > *k_deg_*, slowed export does not affect cytoplasmic noise. However, when *k_exp_* < *k_deg_*, slowed export can cause a decrease in cytoplasmic noise. Data points are mean and error bars are standard deviation of change in cytoplasmic noise across the *possible* parameter regimes for *k_on_*, *k_off_* and *k_tx_*. The shaded area represents ~85% of genes from a previous genome wide study (Bahar Halpern et al., 2015a). **(B)** Predicted change in protein noise (normalized to max) when altering only *k_exp_*. In the regime where *k_exp_* approaches or is < *k_deg_*, simulations predict that for a translation rate of 10 hr^-1^ and a two-hour protein half-life, a decrease in export rate does not propagate to protein noise. **(C)** Nuclear (open) and cytoplasmic (full) mean mRNA count (error bars represent population SEM) and **(D)** ratio of cytoplasmic-to-nuclear noise ratio during treatment with 0.3 (red), 0.6 (blue) and 1 (green) ng/mL of leptomycin B for a range of time periods. mRNA is expressed in an isoclonal populations of Jurkat cells from HIV’s LTR promoter. Observed cytotoxicity marked by the shaded area. **(E)** Calculated rates for all isoclonal populations of Jurkat cells expressing mRNA from HIV’s LTR promoter. **(F)** Calculated export and degradation rates of mRNA expressed from the LTR, in untreated cells (blue), TNF treated cells (yellow) and from fitting an exponential decay to mRNA levels in cells post-Actinomycin D treatment–a transcriptional inhibitor (grey). **(G)** Treatment of Jurkat cells expressing mRNA from the SV40 and UBC promoters with leptomycin B, showed no significant difference in the nuclear mRNA distributions.

**Figure S5.**
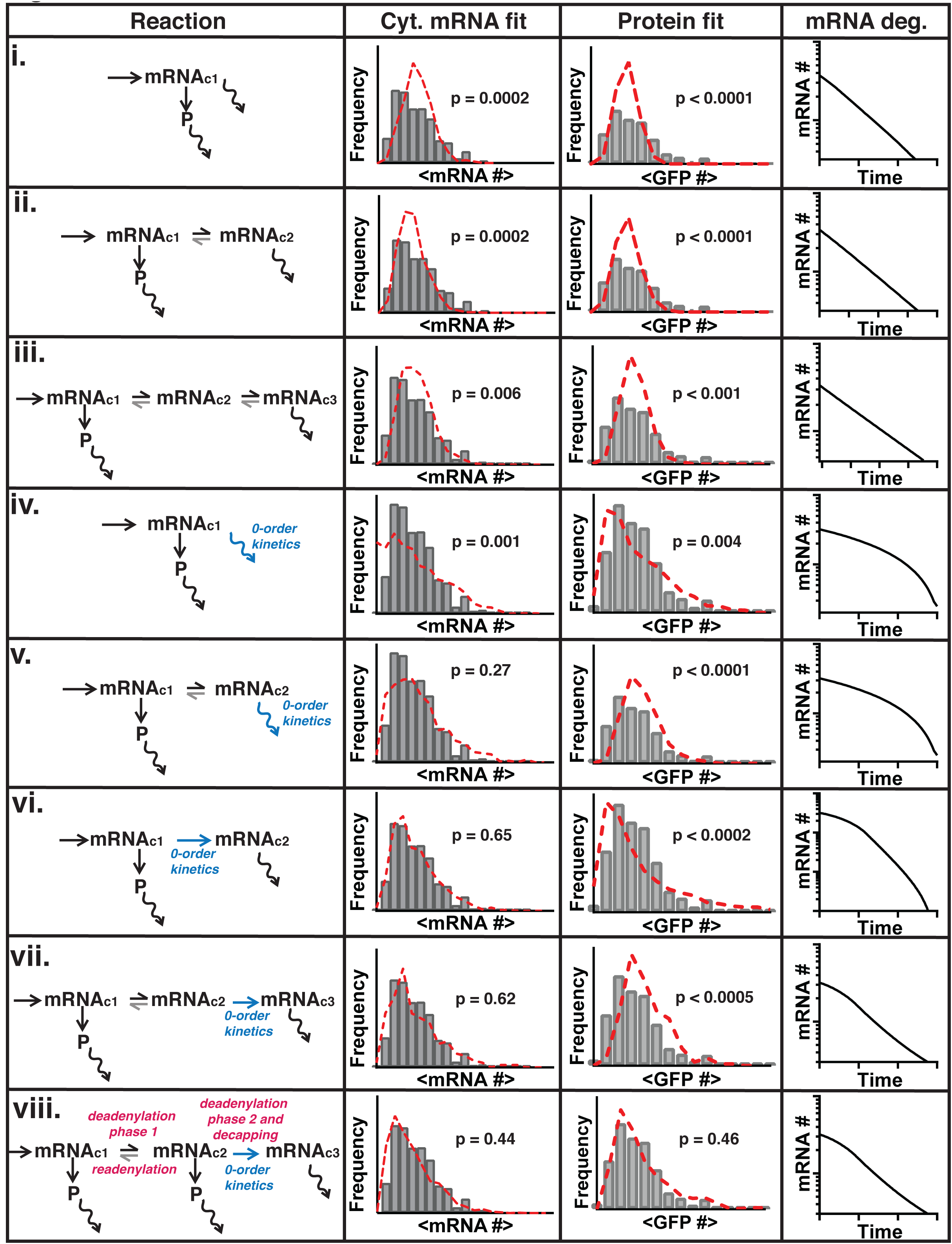
Series of models of increasing complexity compared to experimental mRNA and protein distribution, related to Figure 5. **(i-viii)** Model used (first column) to compare to experimental cytoplasmic mRNA (grey bars, second column) and protein (grey bars, third column) distribution and to model mRNA degradation kinetics (fourth column). **(i-iii)** Single to multiple mRNA states followed by first order mRNA degradation. **(iv-v)** Single to multiple mRNA states with zero-order mRNA degradation (blue arrow). **(vi-viii)** Multiple mRNA states with the rate of mRNA entering the degradation-competent state exhibiting zero-order kinetics (blue arrow), followed by first order mRNA degradation.

**Figure S6.**
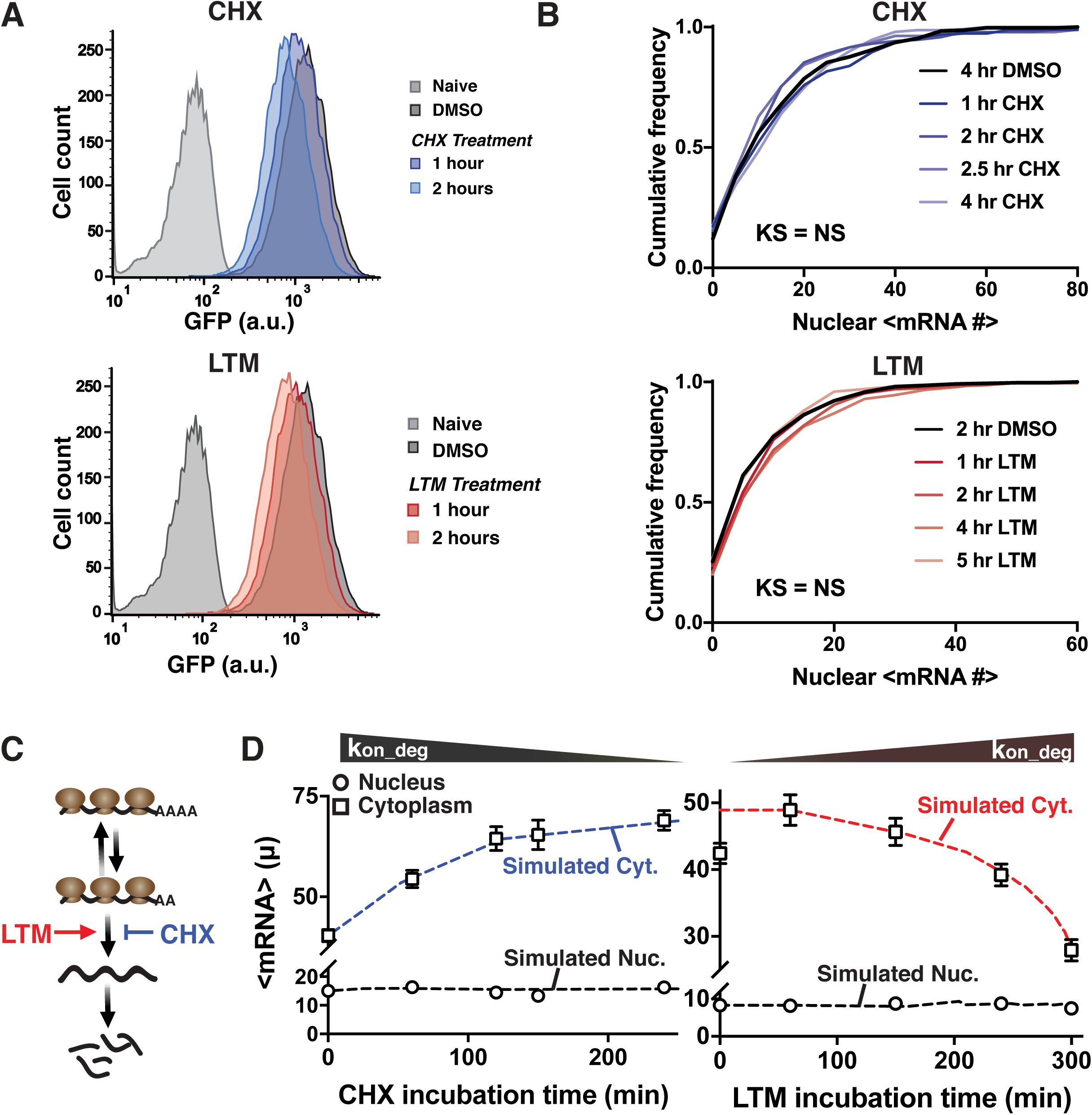
Translation inhibition changes degradation kinetics, related to Figure 5. **(A)** Flow cytometry histogram of human T lymphocytes (Jurkat) expressing d_2_GFP from the LTR promoter (iso. A3), treated with either 100 μM of CHX (top) or 50 μM of LTM (bottom). **(B)** Cumulative frequency distribution of nuclear mRNAs expressed from the HIV-1 LTR promoter (iso. A3), treated with either 100 μM of CHX (top) or 50 μM of LTM (bottom), KS-test is against the DMSO control. **(C)** Lactimidomycin (LTM) and cyclohexamide (CHX) are predicted to have opposite effects on two-state degradation. **(D)** HIV LTR–expressed cytoplasmic mRNA (squares) increases over time in an isoclonal population of human T lymphocytes treated with CHX and decreases over time in cells treated with LTM, while mean nuclear mRNA (circles) remains constant in both cases (black). Simulated nuclear and cytoplasmic means for decreasing (blue) and increasing (red) *k_on_deg_* are shown by the dashed lines.

**Figure S7.**
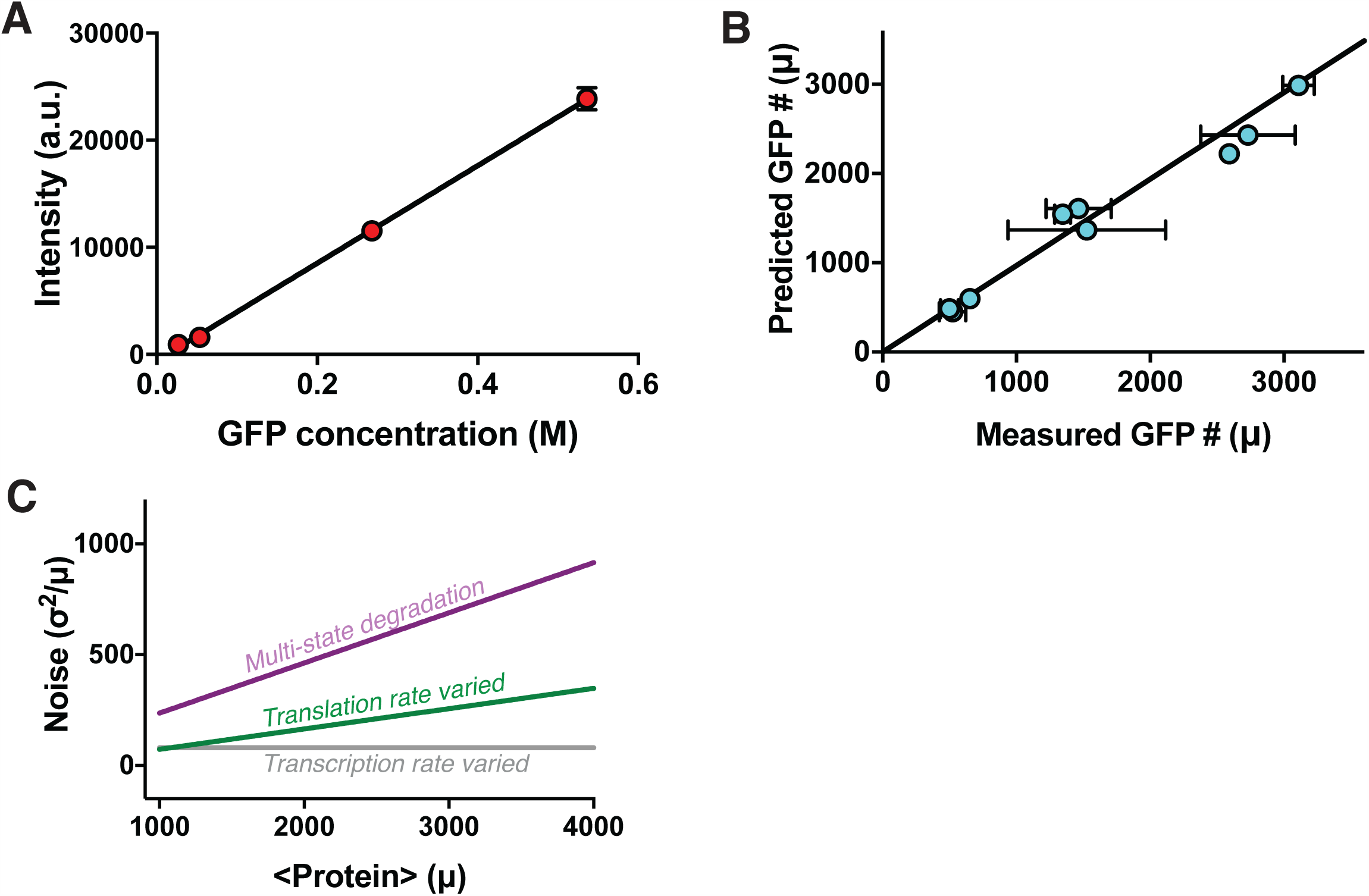
Protein noise correlates with cytoplasmic mRNA noise, related to Figure 6. **(A)** Recombinant eGFP calibration curve for confocal microscope. **(B)** Isoclonal population of Jurkat cells expressing mRNA from HIV’s long terminal repeat (LTR) promoter. Non-Poissonian degradation and translation model, simulated using experimentally derived rate parameters, accurately predicted protein mean. **(C)** Trend in protein noise (σ^2^/μ) versus mean (μ), when only transcription rate is varied (grey), only translation rate is varied (green) and multi-state degradation is introduced (purple).

**Figure S8.**
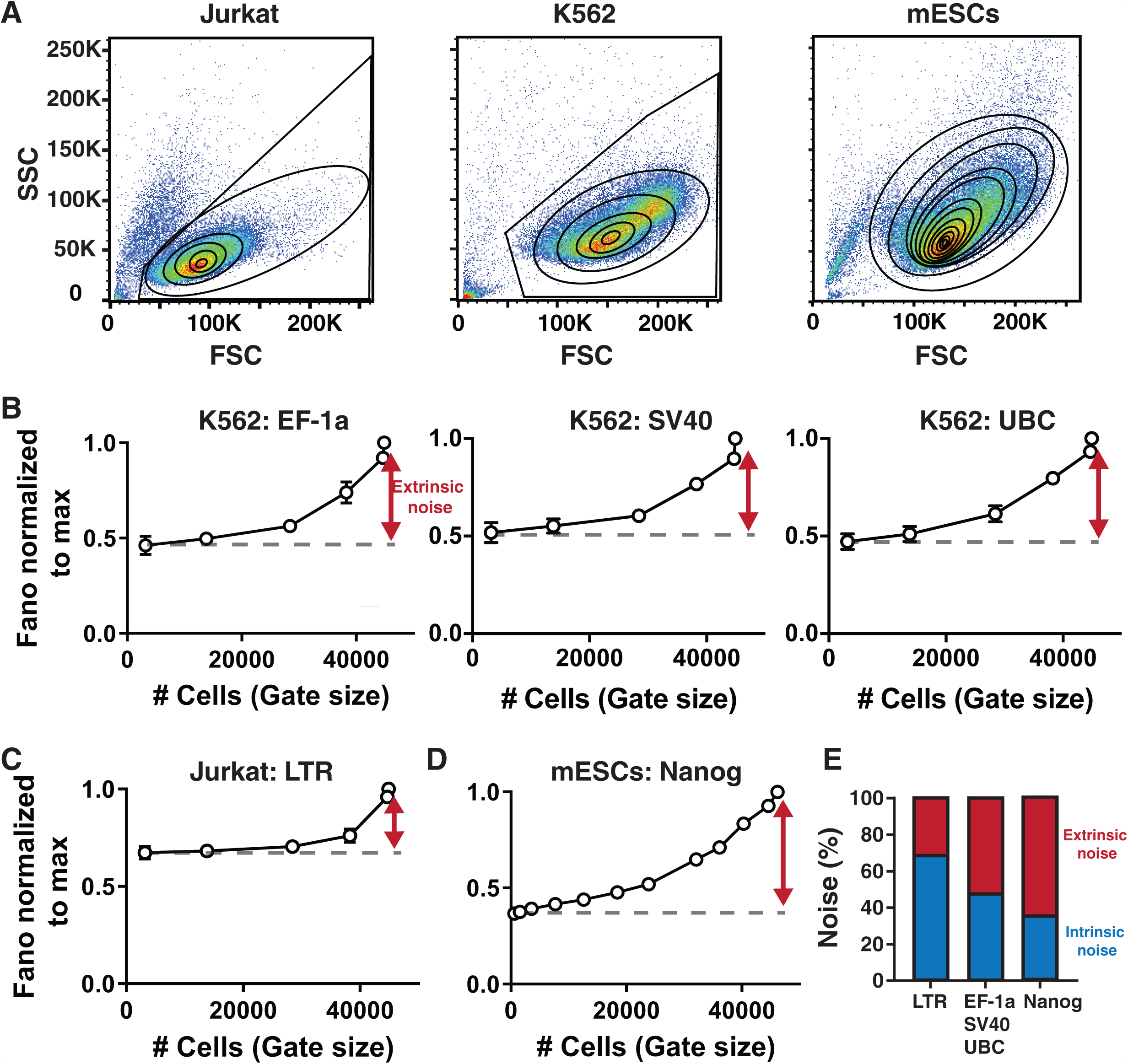
Extrinsic noise contributions to total cell-cell variability, related to Figure 6. **(A)** Representative flow cytometry dot plot showing the different sized gates used for the extrinsic noise analysis for human T lymphocytes (Jurkat), human myeloid leukemia cells (K562) and mouse embryonic stem cells (mESCs). **(B-D)** Noise (σ^2^/μ) normalized to max versus number of cells in the respective gates from (A) d_2_GFP expressed from: **(C)** the UBC and EF-1α promoters, and GFP expressed from the SV40 promoter; **(C)** the LTR promoter and; **(D)** from the NANOG promoter. **(E)** Contributions from extrinsic (red) and intrinsic (blue) noise towards total noise for 5 different promoters.

## REFERENCES

Arias, A.M., and Hayward, P. (2006). Filtering transcriptional noise during development: concepts and mechanisms. Nat Rev Genet 7, 34–44.

Bahar Halpern, K., Caspi, I., Lemze, D., Levy, M., Landen, S., Elinav, E., Ulitsky, I., and Itzkovitz, S. (2015). Nuclear Retention of mRNA in Mammalian Tissues. Cell Reports 13, 2653–2662.

Balázsi, G., van Oudenaarden, A., and Collins, James J. (2011). Cellular Decision Making and Biological Noise: From Microbes to Mammals. Cell 144, 910–925.

Bar-Even, A., Paulsson, J., Maheshri, N., Carmi, M., O’Shea, E., Pilpel, Y., and Barkai, N. (2006). Noise in protein expression scales with natural protein abundance. Nat Genet 38, 636–643.

Bartel, D.P. (2004). MicroRNAs: Genomics, Biogenesis, Mechanism, and Function. Cell 116, 281–297.

Battich, N., Stoeger, T., and Pelkmans, L. (2015). Control of Transcript Variability in Single Mammalian Cells. Cell 163, 1596–1610.

Beaumont, H.J.E., Gallie, J., Kost, C., Ferguson, G.C., and Rainey, P.B. (2009). Experimental evolution of bet hedging. Nature 462, 90–93.

Becskei, A., Kaufmann, B.B., and van Oudenaarden, A. (2005). Contributions of low molecule number and chromosomal positioning to stochastic gene expression. Nat Genet 37, 937–944.

Bensaude, O. (2011). Inhibiting eukaryotic transcription: Which compound to choose? How to evaluate its activity? Transcription 2, 103–108.

Blake, W.J., Balázsi, G., Kohanski, M.A., Isaacs, F.J., Murphy, K.F., Kuang, Y., Cantor, C.R., Walt, D.R., and Collins, J.J. (2006). Phenotypic Consequences of Promoter-Mediated Transcriptional Noise. Molecular Cell 24, 853–865.

Blake, W.J., Kaern, M., Cantor, C.R., and Collins, J.J. (2003). Noise in eukaryotic gene expression. Nature 422, 633–637.

Boireau, S., Maiuri, P., Basyuk, E., de la Mata, M., Knezevich, A., Pradet-Balade, B., Bäcker, V., Kornblihtt, A., Marcello, A., and Bertrand, E. (2007). The transcriptional cycle of HIV-1 in real-time and live cells. J Cell Biol 179, 291–304.

Casolari, J.M., Brown, C.R., Komili, S., West, J., Hieronymus, H., and Silver, P.A. (2004). Genome-Wide Localization of the Nuclear Transport Machinery Couples Transcriptional Status and Nuclear Organization. Cell 117, 427–439.

Choubey, S., Kondev, J., and Sanchez, A. (2015). Deciphering Transcriptional Dynamics In Vivo by Counting Nascent RNA Molecules. PLOS Computational Biology 11, e1004345.

Coffman, V.C., and Wu, J.-Q. (2014). Every laboratory with a fluorescence microscope should consider counting molecules. Molecular Biology of the Cell 25, 1545–1548.

Dar, R.D., Hosmane, N.N., Arkin, M.R., Siliciano, R.F., and Weinberger, L.S. (2014). Screening for noise in gene expression identifies drug synergies. Science 344, 1392–1396.

Dar, R.D., Razooky, B.S., Singh, A., Trimeloni, T.V., McCollum, J.M., Cox, C.D., Simpson, M.L., and Weinberger, L.S. (2012). Transcriptional burst frequency and burst size are equally modulated across the human genome. Proceedings of the National Academy of Sciences 109, 17454–17459.

Decker, C.J., and Parker, R. (2012). P-Bodies and Stress Granules: Possible Roles in the Control of Translation and mRNA Degradation. Cold Spring Harbor Perspectives in Biology 4.

Duh, E.J., Maury, W.J., Folks, T.M., Fauci, A.S., and Rabson, A.B. (1989). Tumor necrosis factor alpha activates human immunodeficiency virus type 1 through induction of nuclear factor binding to the NF-kappa B sites in the long terminal repeat. Proceedings of the National Academy of Sciences 86, 5974–5978.

Felber, B.K., Hadzopoulou-Cladaras, M., Cladaras, C., Copeland, T., and Pavlakis, G.N. (1989). rev protein of human immunodeficiency virus type 1 affects the stability and transport of the viral mRNA. Proc Natl Acad Sci U S A 86, 1495–1499.

Fraser, H.B., Hirsh, A.E., Giaever, G., Kumm, J., and Eisen, M.B. (2004). Noise minimization in eukaryotic gene expression. PLoS Biol 2, e137.

Garg, S., and Sharp, P.A. (2016). Single-cell variability guided by microRNAs. Science 352, 1390.

Garneau, N.L., Wilusz, J., and Wilusz, C.J. (2007). The highways and byways of mRNA decay. Nat Rev Mol Cell Biol 8, 113–126.

Gilbert, Luke A., Larson, Matthew H., Morsut, L., Liu, Z., Brar, Gloria A., Torres, Sandra E., Stern-Ginossar, N., Brandman, O., Whitehead, Evan H., Doudna, Jennifer A., et al. (2013). CRISPR-Mediated Modular RNA-Guided Regulation of Transcription in Eukaryotes. Cell 154, 442–451.

Gillespie, D.T. (1977). Exact stochastic simulation of coupled chemical reactions. The Journal of Physical Chemistry 81, 2340–2361.

Hao, S., and Baltimore, D. (2013). RNA splicing regulates the temporal order of TNF-induced gene expression. Proceedings of the National Academy of Sciences of the United States of America 110, 11934–11939.

Hasty, J., McMillen, D., and Collins, J.J. (2002). Engineered gene circuits. Nature 420, 224–230.

Horvathova, I., Voigt, F., Kotrys, A.V., Zhan, Y., Artus-Revel, C.G., Eglinger, J., Stadler, M.B., Giorgetti, L., and Chao, J.A. (2017). The Dynamics of mRNA Turnover Revealed by Single-Molecule Imaging in Single Cells. Molecular Cell 68, 615–625.e619.

Kaern, M., Elston, T.C., Blake, W.J., and Collins, J.J. (2005). Stochasticity in gene expression: from theories to phenotypes. Nat Rev Genet 6, 451–464.

Kepler, T.B., and Elston, T.C. (2001). Stochasticity in transcriptional regulation: origins, consequences, and mathematical representations. Biophys J 81, 3116–3136.

Kim, D.W., Uetsuki, T., Kaziro, Y., Yamaguchi, N., and Sugano, S. (1990). Use of the Human Elongation Factor-1-Alpha Promoter as a Versatile and Efficient Expression System. Gene 91, 217–223.

LaGrandeur, T., and Parker, R. (1999). The cis acting sequences responsible for the differential decay of the unstable MFA2 and stable PGK1 transcripts in yeast include the context of the translational start codon. RNA 5, 420–433.

Lee, S., Liu, B., Lee, S., Huang, S.-X., Shen, B., and Qian, S.-B. (2012). Global mapping of translation initiation sites in mammalian cells at single-nucleotide resolution. Proceedings of the National Academy of Sciences 109, E2424–E2432.

Malim, M.H., Hauber, J., Fenrick, R., and Cullen, B.R. (1988). Immunodeficiency virus rev trans-activator modulates the expression of the viral regulatory genes. Nature 335, 181–183.

Mittler, J.E., Sulzer, B., Neumann, A.U., and Perelson, A.S. (1998). Influence of delayed viral production on viral dynamics in HIV-1 infected patients. Mathematical Biosciences 152, 143–163.

Munsky, B., Neuert, G., and van Oudenaarden, A. (2012). Using Gene Expression Noise to Understand Gene Regulation. Science 336, 183–187.

Newman, J.R.S., Ghaemmaghami, S., Ihmels, J., Breslow, D.K., Noble, M., DeRisi, J.L., and Weissman, J.S. (2006). Single-cell proteomic analysis of S. cerevisiae reveals the architecture of biological noise. Nature 441, 840–846.

Ossareh-Nazari, B., Bachelerie, F., and Dargemont, C. (1997). Evidence for a role of CRM1 in signal-mediated nuclear protein export. Science 278, 141–144.

Ozbudak, E.M., Thattai, M., Kurtser, I., Grossman, A.D., and van Oudenaarden, A. (2002). Regulation of noise in the expression of a single gene. Nat Genet 31, 69–73.

Padovan-Merhar, O., Nair, Gautham P., Biaesch, Andrew G., Mayer, A., Scarfone, S., Foley, Shawn W., Wu, Angela R., Churchman, L.S., Singh, A., and Raj, A. (2015). Single Mammalian Cells Compensate for Differences in Cellular Volume and DNA Copy Number through Independent Global Transcriptional Mechanisms. Molecular Cell 58, 339–352.

Parker, R. (2012). RNA Degradation in *Saccharomyces cerevisae*. Genetics 191, 671–702.

Pelechano, V., Wei, W., and Steinmetz, Lars M. (2015). Widespread Co-translational RNA Decay Reveals Ribosome Dynamics. Cell 161, 1400–1412.

Raj, A., Peskin, C.S., Tranchina, D., Vargas, D.Y., and Tyagi, S. (2006). Stochastic mRNA Synthesis in Mammalian Cells. PLOS Biology 4, e309.

Raj, A., and van Oudenaarden, A. (2008). Nature, Nurture, or Chance: Stochastic Gene Expression and Its Consequences. Cell 135, 216–226.

Rouzine, I.M., Weinberger, A.D., and Weinberger, L.S. (2015). An evolutionary role for HIV latency in enhancing viral transmission. Cell 160, 1002–1012.

Sanchez, A., and Golding, I. (2013). Genetic Determinants and Cellular Constraints in Noisy Gene Expression. Science 342, 1188–1193.

Schmiedel, J.M., Klemm, S.L., Zheng, Y., Sahay, A., Blüthgen, N., Marks, D.S., and van Oudenaarden, A. (2015). MicroRNA control of protein expression noise. Science 348, 128.

Simpson, M.L., Cox, C.D., and Sayler, G.S. (2003). Frequency domain analysis of noise in autoregulated gene circuits. Proceedings of the National Academy of Sciences 100, 4551–4556.

Singh, A., and Bokes, P. (2012). Consequences of mRNA Transport on Stochastic Variability in Protein Levels. Biophysical Journal 103, 1087–1096.

Singh, A., Razooky, B., Cox, C.D., Simpson, M.L., and Weinberger, L.S. (2010). Transcriptional Bursting from the HIV-1 Promoter Is a Significant Source of Stochastic Noise in HIV-1 Gene Expression. Biophysical Journal 98, L32–L34.

Thattai, M., and van Oudenaarden, A. (2001). Intrinsic noise in gene regulatory networks. Proceedings of the National Academy of Sciences 98, 8614–8619.

Tilgner, H., Knowles, D.G., Johnson, R., Davis, C.A., Chakrabortty, S., Djebali, S., Curado, J., Snyder, M., Gingeras, T.R., and Guigó, R. (2012). Deep sequencing of subcellular RNA fractions shows splicing to be predominantly co-transcriptional in the human genome but inefficient for lncRNAs. Genome Research 22, 1616–1625.

Urcuqui-Inchima, S., Patiño, C., Zapata, X., García, M.P., Arteaga, J., Chamot, C., Kumar, A., and Hernandez-Verdun, D. (2011). Production of HIV Particles Is Regulated by Altering Sub-Cellular Localization and Dynamics of Rev Induced by Double-Strand RNA Binding Protein. PLOS ONE 6, e16686.

Watanabe, M., Fukuda, M., Yoshida, M., Yanagida, M., and Nishida, E. (1999). Involvement of CRM1, a nuclear export receptor, in mRNA export in mammalian cells and fission yeast. Genes to Cells 4, 291–297.

Weinberger, L.S., Burnett, J.C., Toettcher, J.E., Arkin, A.P., and Schaffer, D.V. (2005). Stochastic gene expression in a lentiviral positive-feedback loop: HIV-1 Tat fluctuations drive phenotypic diversity. Cell 122, 169–182.

Wolf, L., Silander, O.K., and van Nimwegen, E. (2015). Expression noise facilitates the evolution of gene regulation. eLife 4, e05856.

Xiong, L.-p., Ma, Y.-q., and Tang, L.-h. (2010). Attenuation of transcriptional bursting in mRNA transport. Physical Biology 7, 016005.

Yamashita, A., Chang, T.-C., Yamashita, Y., Zhu, W., Zhong, Z., Chen, C.-Y.A., and Shyu, A.-B. (2005). Concerted action of poly(A) nucleases and decapping enzyme in mammalian mRNA turnover. Nature Structural & Molecular Biology 12, 1054.

